# The adaptive potential of the middle domain of yeast Hsp90

**DOI:** 10.1101/832022

**Authors:** Pamela A. Cote-Hammarlof, Inês Fragata, Julia Flynn, David Mavor, Konstantin B. Zeldovich, Claudia Bank, Daniel N.A. Bolon

**Author notes:** Co-corresponding authors Daniel N.A. Bolon, Claudia Bank. Co-first authors (Pamela A. Cote-Hammarlof, Inês Fragata). Centre for Ecology and Evolution, Faculdade de Ciências da Universidade de Lisboa, Portugal (Inês Fragata).

## Abstract

The distribution of fitness effects (DFE) of new mutations across different environments quantifies the potential for adaptation in a given environment and its cost in others. So far, results regarding the cost of adaptation across environments have been mixed, and most studies have sampled random mutations across different genes. Here, we quantify systematically how costs of adaptation vary along a large stretch of protein sequence by studying the DFEs of the same ≈2300 amino-acid changing mutations obtained from deep mutational scanning of 119 amino acids in the middle domain of the heat-shock protein Hsp90 in five environments. This region is known to be important for client binding, stabilization of the Hsp90 dimer, stabilization of the N-terminal-Middle and Middle-C-terminal interdomains, and regulation of ATPase-chaperone activity. Interestingly, we find that fitness correlates well across diverse stressful environments, with the exception of one environment, diamide. Consistent with this result, we find little cost of adaptation; on average only one in seven beneficial mutations is deleterious in another environment. We identify a hotspot of beneficial mutations in a region of the protein that is located within an allosteric center. The identified protein regions that are enriched in beneficial, deleterious, and costly mutations coincide with residues that are involved in the stabilization of Hsp90 interdomains and stabilization of client binding interfaces, or residues that are involved in ATPase chaperone activity of Hsp90. Thus, our study yields information regarding the role and adaptive potential of a protein sequence that complements and extends known structural information.

## Introduction

The distribution of fitness effects (DFE) determines the proportions of new mutations that are beneficial, deleterious or neutral (Eyre-Walker & Keightley 2007, Loewe 2010, Bataillon & Bailey 2014). It provides a snapshot of the robustness of the genome to changes in the DNA and carries information about the expected amount of genetic diversity within populations. Moreover, the beneficial part of the DFE informs on the adaptive potential of populations when introduced into a new environment (Sniegowski & Gerrish 2010, Bataillon & Bailey 2014). However, beneficial mutations in one environment can be deleterious in another, potentially resulting in so-called costs of adaptation (Bataillon et al 2011), also termed antagonistic pleiotropy. So far, it has been difficult to address the prevalence of costs of adaptation, because measuring the fitness of the same mutations across various environments is not straightforward. Specifically, previous studies using a selection of mutations obtained from laboratory evolution or mutation accumulation experiments found that antagonistic pleiotropy was rare (e.g., Ostrowski et al 2005, Dillon et al 2016, Sane et al 2018). It is unknown whether this pattern holds for an unbiased selection of mutants.

Comparing the fitness of a large unbiased selection of mutants across environments has become feasible with the advancement of deep mutational scanning. Developed around a decade ago, deep mutational scanning allows for the assessment of the complete DFE of a focal genomic region in some genetically modifiable and fast-growing model species, using a combination of site-directed mutagenesis and deep sequencing (Fowler et al. 2010, Hietpas et al 2011, 2012, Boucher et al. 2014, Logacheva et al 2016). Deep mutational scanning studies from single environments usually report a bimodal DFE with two peaks that represent neutral and strongly deleterious mutations, respectively (Acevedo et al 2014, Hietpas et al. 2011, Boucher et al 2014). Despite the ample use of this approach (Melamed et al 2013, Boucher et al 2014, Doud et al 2016, Sarkisyan et al 2016), few studies have quantified the impact of environmental challenges on the shape of these DFEs (but see Hietpas et al 2013, Bank et al., 2014). However, experimentally quantifying the DFE across environments is important because natural environments are constantly changing, which can have diverse evolutionary consequences that are directly related to the shape of the DFE and the costs of adaptation (Dhar et al. 2011, Arribas et al. 2014, Mumby and van Woesik 2014, Brennan et al. 2017).

The consequences of environmental challenges on organisms manifest at many biological levels, including the protein level. For example, increased temperature can cause protein unfolding and aggregation, which can disrupt function and ultimately affect organismal fitness and survival (Richter et al. 2010). Chaperones, such as the heat shock protein Hsp90, help the cellular machinery survive stress conditions (Chen et al. 2018). By buffering deleterious fitness effects in stress conditions, chaperones are important for the short-term response to new and recurring environmental challenges. On a longer time scale, their buffering effect has also been argued to facilitate the maintenance of standing genetic variation elsewhere in the genome that can enable rapid genetic adaptation to new stress conditions (Rutherford 2003, Barrett and Schluter 2008, Jarosz and Lindquist 2010, Fitzgerald and Rosenberg 2019). Therefore, it is important to understand how chaperones evolve, and how the selection pressure on a chaperone changes upon exposure to different environmental challenges.

The heat shock protein Hsp90 is a chaperone that plays an essential role in protecting cells from environmental stress and that is required at elevated levels for yeast growth at high temperature (Borkovich et al. 1989). Recent studies using systematic mutagenesis have begun to address how mutations to a strongly conserved client binding site of Hsp90 can impact evolutionary adaptation in yeast (Hietpas et al. 2013). Multiple mutations in a nine amino acid client-binding site of yeast Hsp90 provided a growth advantage under elevated salinity conditions (Hietpas et al. 2013, Bank et al 2014). A recent larger-scale study found that at low Hsp90 expression levels, changes in environment greatly changed the shape of the DFE, and that some environments showed a higher prevalence of both beneficial and deleterious mutations (Flynn et al 2020). However, another previous study proposed that at low expression the fitness effects of mutations, especially deleterious ones, should be larger (Jiang et al 2013). Thus, it is unknown how much of the observed effect was due to expression level and how much was due to environmental changes.

Here we examined a 119 amino acid region (encompassing positions 291-409, Fig. 1) of the middle domain of yeast Hsp90 (aka Hsp82) at normal expression levels. Several studies have demonstrated that the middle domain of Hsp90 plays a prominent role in client binding (Nathan and Lindquist 1995, Nathan et al. 1997, Meyer et al. 2003, Hawle et al. 2006, Hagn et al. 2011), and suggested that mutations in this region may impact the relative affinity or priority of different clients and physiological pathways with the potential to provide an adaptive benefit (Hagn et al. 2011, Zhang et al. 2005, Sato et al. 2000, Karagoz et al. 2014, 2015, Verba et al. 2016, Czemeres et al. 2017).

**Figure 1:**
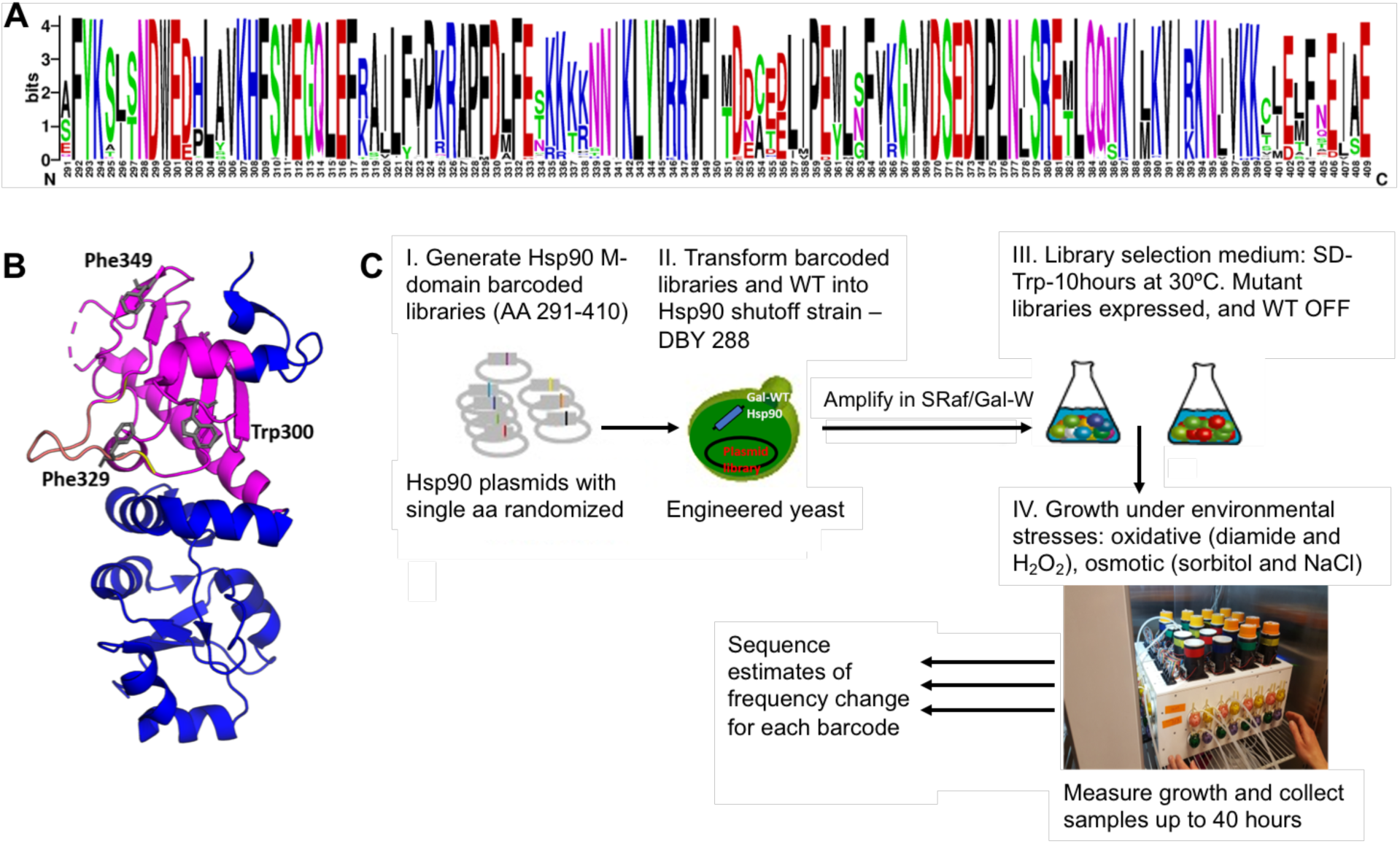
Middle domain of Hsp90. (A) Amino acid conservation of yeast Hsp90 compared to 261 eukaryotic sequences. Relative height of the amino acid indicates degree of conservation. (B) Structural representation based on 1HK7.PDB of the middle domain of Hsp90, with amino acids 291-409 that are the focus of this study highlighted in purple. A solvent exposed amphipathic loop implicated in client binding (amino acids 327-341) is highlighted in yellow and residues implicated in client binding (amino acids W300, F329, F349) are shown as gray sticks. (C) Systematic approach to measure the adaptive potential of the middle domain of yeast Hsp90 (amino acids 291-409).

We used the EMPIRIC approach of deep mutational scanning (Hietpas et al 2011) to estimate the selection coefficients of all amino-acid changing mutations in the middle domain of yeast Hsp90 across five environments. Using the inferred DFEs, we identified regions of the middle domain that stood out with respect to their potential for adaptation upon environmental change. Moreover, as some environments share the type of stress that they induce, namely osmotic stress (0.5M salinity or 0.6M sorbitol) and oxidative stress (0.6 mM H202 or 0.85mM diamide), we were able to study the impact of the type of stress on the adaptive potential of new mutations. To this end we: a) quantified the impact of environmental changes on the overall shape of the DFE, b) identified regions that showed the largest proportions of beneficial or deleterious mutations, respectively, c) quantified hotspots of costs of adaptation, and d) compared the identified regions with known client binding or other structurally important sites to connect the phenotypic and fitness effects of mutations. Altogether, we mapped potential protein regions that may play an important role in adaptation to different environments.

## Results and Discussion

To investigate the adaptive potential of the middle domain of yeast Hsp90 we used systematic site-directed mutagenesis of amino acid positions 291-409 that include known client binding sites. This resulted in ≈2300 amino-acid changing mutations, For which selection coefficients were estimated from 2-3 replicates of bulk competitions that were performed in five environments. We focused our analysis on standard lab conditions and four environmental stresses that affect growth rate (Supplementary figure 1) in yeast (Gasch et al. 2000).

### DFEs across all environments show many wild-type like mutations

The shape of the DFE indicates the relative importance of purifying or directional selection as compared with neutral evolution. In apparent contrast to the strong conservation of the Hsp90 middle domain in natural populations, we observed DFEs with mostly wild-type like mutations across all environments (Supplementary Figure 2, 3). Throughout the manuscript, we use the term “wild-type like” to denote mutations that are indistinguishable from the wild-type reference in the limit of experimental accuracy, see Material and Methods. We categorized mutants as wild-type like if the 95% confidence interval of the estimated selection coefficient overlapped with 0 (see also Supplementary Table 1, Materials & Methods). According to this criterion, between 50% (in the H_2_0_2_ environment) and 65% (in the standard environment) of mutations showed a fitness effect that is indistinguishable from the reference type. Large numbers of wild-type like mutations have been observed previously in deep mutational scanning studies of DFEs (Soskine and Tawfik 2010, Hietpas et al 2013, Melamed et al 2013, Bank et al 2014, Hom et al 2019). Both biological and technical factors can be invoked as an explanation for the large number of wild-type like mutations. Firstly, the resolution of the experiment is likely much lower than the resolution at which natural selection may act in large yeast populations (Ohta 1992, Boucher et al. 2016). Secondly, selection pressures in the laboratory might differ greatly from those in nature (e.g., Reznick & Ghalambor 2005, Kvitek and Sherlock 2013). Finally and relatedly, natural environments might be fundamentally different and rapidly fluctuating (Mustonen and Lässig 2009). For example, a recent study of the DFE of the full Hsp90 sequence found that mutations which were tolerated across a set of diverse environments were those most likely observed in natural sequences (Flynn et al., 2020). We repeated the analysis of Flynn et al. 2020 and found the same pattern: our estimated DFEs across all environments of the subset of variants observed in nature (amino acid mutations observed across 261 eukaryotic sequences, see Material and Methods) showed an elevated peak around 0. This indicates a further enrichment of wild-type like mutations in the subset of naturally observed variants as compared with the full data set (Supplementary Figure 5).

**Figure 2:**
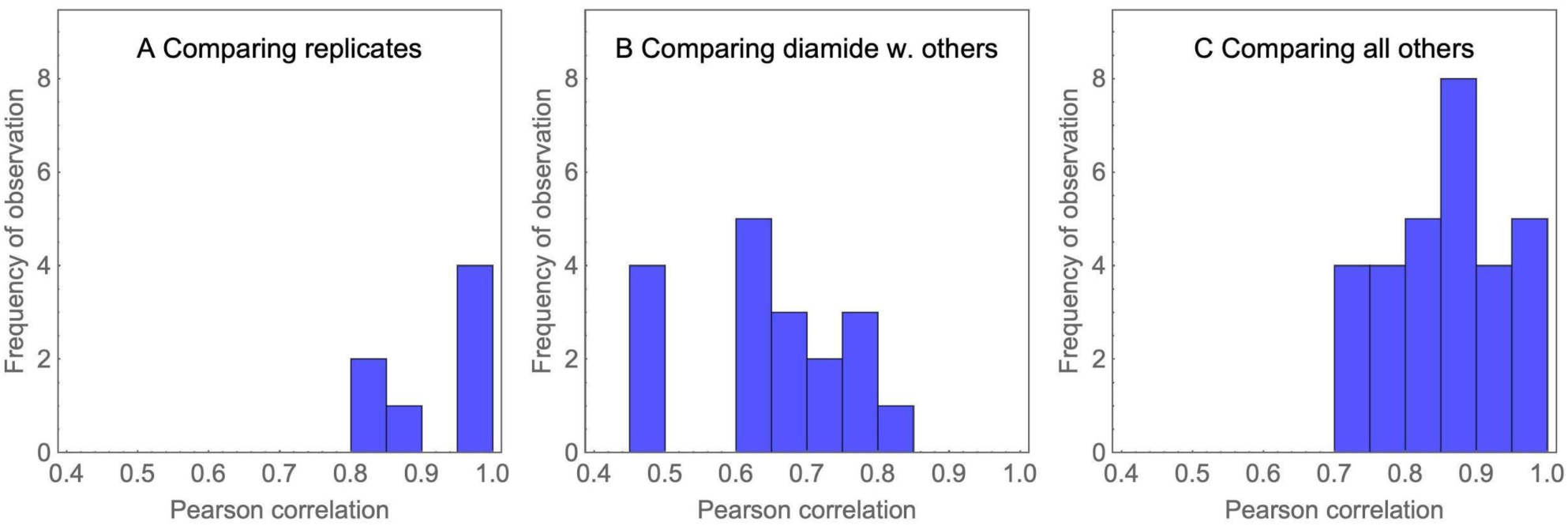
Histogram of correlations of fitness effects among replicates and environments reveals that diamide stands out as different. The histograms correspond to comparisons of replicates (A), comparisons that include diamide (B), and all other comparisons (C). Correlations between replicates and between pairs of environments are in general high, whereas correlations between diamide and the other environments tend to be lower. Altogether, this suggests that cells react in unique ways to the oxidative stress enacted by exposure to diamide, and that Hsp90 plays a specific role in the response to this stress.

**Figure 3:**
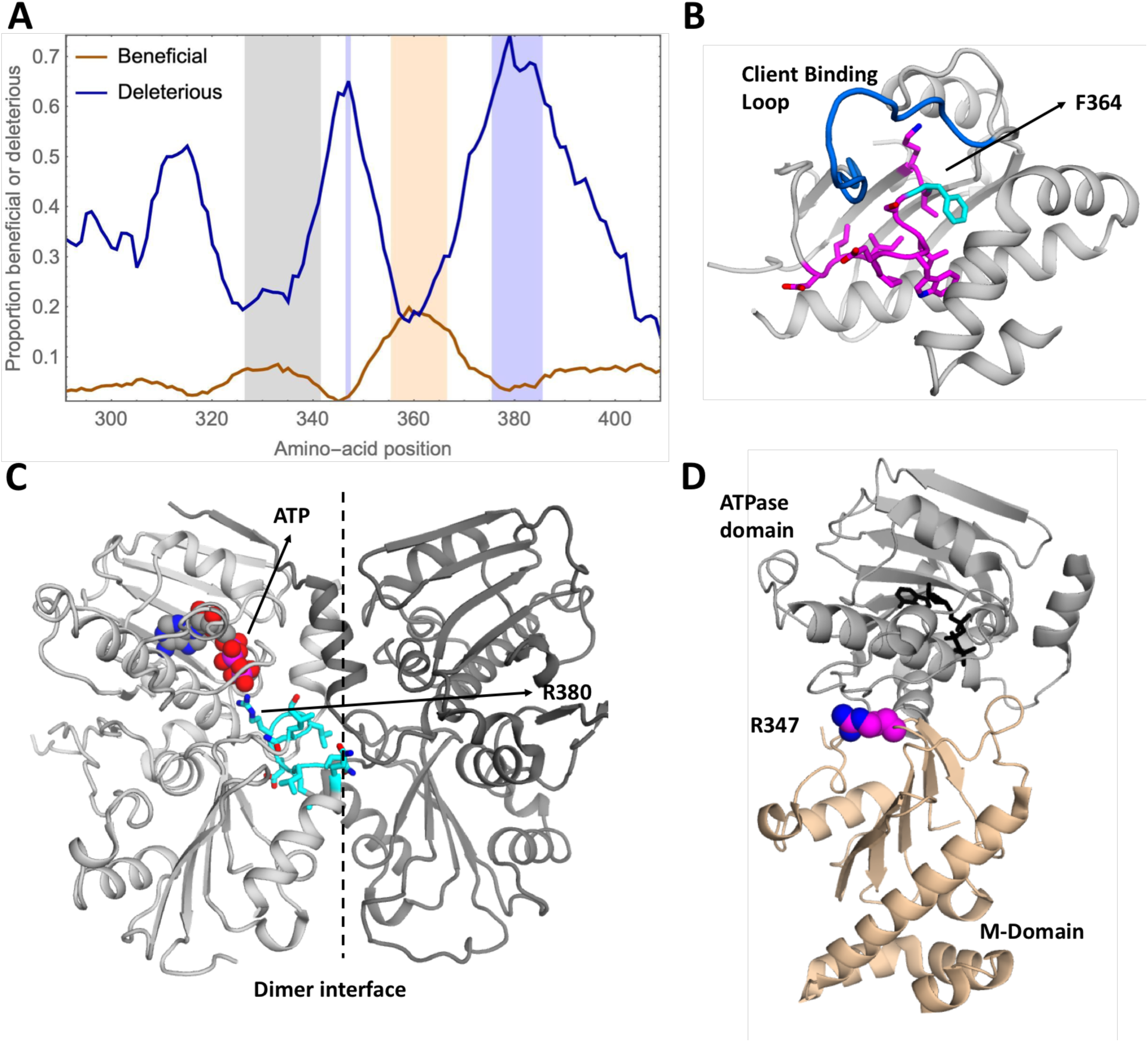
Hotspots of beneficial and deleterious mutations in the middle domain do not coincide with known client binding loop. A) Average proportion of beneficial and deleterious mutations along the genome across all environments, illustrated using a 10-amino-acid sliding window. The region with the 10% largest proportions of beneficial mutations is highlighted in light orange (position 356-366), the region with the 10% largest proportions of deleterious mutations is highlighted in light blue (positions 347 and 376-385). A known client binding loop (positions 327-341) is highlighted in light gray. To avoid biases, the analysis was restricted to subsets of two replicates of all environments. B) Structural representation of the Hsp90 middle domain (1HK7.PDB) illustrating the beneficial hotspot in magenta with residue 364 highlighted in cyan. Biochemical, structural and mutational studies identified positions 327-341 as a client binding loop, which is shown in dark blue. The beneficial hotspot that was identified in our analyses is adjacent to the above mentioned client binding loop. C) Structural representation of the ATPase and middle domains of Hsp90 from 2CG9.PDB. The deleterious hotspot (in cyan) contains a catalytic amino acid and is located at a dimerization interface. It includes residue R380 that stabilizes the leaving phosphate during ATP hydrolysis. Subunits are distinguished with different shading. D) The deleterious hotspot at position 347 is also located at the interface of the ATPase domain and the middle domain and is highlighted in pink and blue.

**Figure 4:**
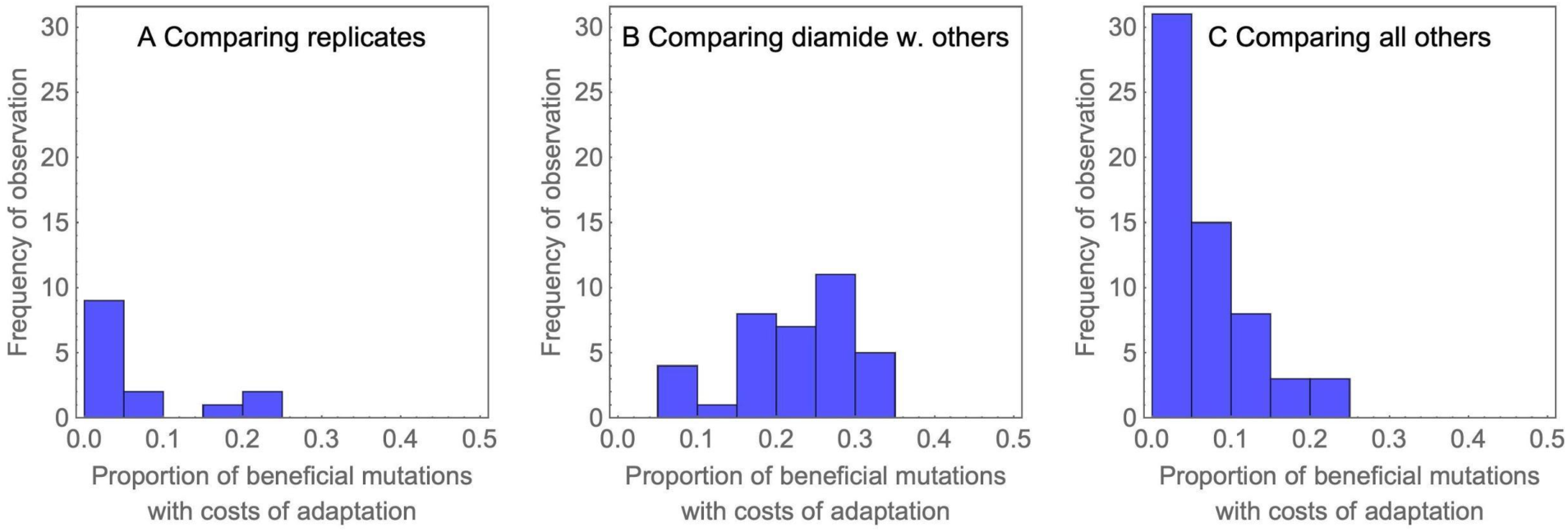
Histogram of proportions of mutations that display costs of adaptation illustrates that costs of adaptation are generally rare, and most likely if comparisons involve diamide environments. The histograms correspond to comparisons of replicates (A), the subset of other comparisons that include diamide (B), and all other comparisons (C).

**Figure 5:**
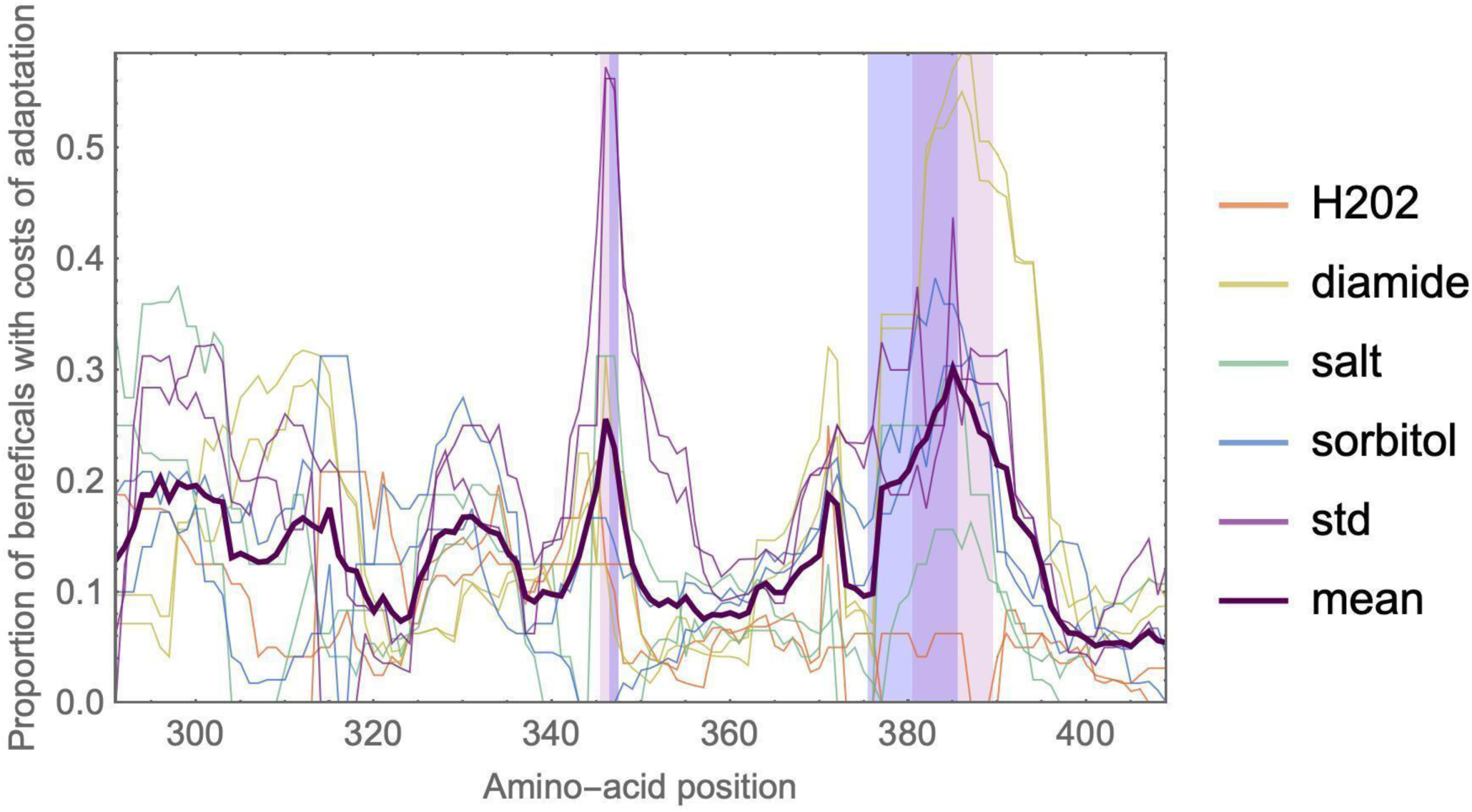
A hotspot of costs of adaptation is located at positions 381-391. The 10% positions with the largest mean proportion of mutations that display costs of adaptation (dark purple line) are highlighted in light purple. This region greatly overlaps with the hotspots of deleterious mutations from Figure 3 (highlighted in blue). Thin curves indicate the mean proportion of beneficial mutations in a focal environment that is deleterious in another environment. A 10-amino-acid sliding window was used to locate and display region-specific effects. To avoid biases, the analysis was restricted to subsets of two replicates of all environments.

Our approach provides a dense scan of the local fitness landscape by measuring the selection coefficient of all amino-acid changing mutations that are available in a single mutational step from the ancestral state. The presence of a large number of wild-type like mutations in various different environments suggests that the local fitness landscape is rather flat, which is at odds with the strong conservation of the protein in yeast. However, further away from the wild type, epistatic interactions may condition the following mutational steps and thereby change the configuration of the fitness landscape, creating fewer attainable mutational paths and constraining evolution on longer time scales (e.g. Weinreich et al 2006, Kryazhimsky et al 2014).

### Correlations across conditions reveal that diamide stands out with respect to mutational effects

We next quantified the correlations of fitness effects across replicates and environments (Figure 2, Supplementary Table 2). Consistent with the high accuracy of the experiment, we observed strong correlations of estimated mutational fitness effects between replicates (Supplementary Figure 6, mean Pearson correlation r=0.91). Across pairs of environments, we observed a large variation of correlations ranging from r=0.48 between diamide and salt to r=0.98 between H_2_0_2_ and the standard environment. In this analysis, diamide clearly stood out by showing consistently lower correlations of fitness effects with other environments than all others (mean correlation including diamide r=0.65, mean correlation of all others r=0.86). This could be because diamide exerts multiple negative effects on the yeast cells. For example, among the environments we investigated, diamide was the only condition that induces the expression of cell wall biosynthesis genes and genes involved in protein secretion and processing in the Endoplasmic Reticulum, which indicates its role in cell wall damage (Gasch et al. 2000). Diamide was also shown to affect individual transcription factors differently from H_2_0_2_, the other oxidative environment in our selection. One example that has been described in the literature is the Yap1 transcription factor, which is nuclear localized and active in diamide, but cytoplasmic and inactive in H_2_0_2_ (Gulshan et al. 2011). Interactions of Hsp90 with other genes may also contribute to fitness differences of the mutations between the tested environmental conditions and mutants that we tested. Further global experimental analyses beyond the scope of this work will be required to determine the molecular features of diamide stress that elicit distinct selection on Hsp90 sequence. Previous studies have reported that mutations had similar fitness effects across environments that shared metabolic features (Ostrowski et al 2005, Dillon et al 2015, Sane et al 2018). Thus, we expected to see stronger correlations of fitness effects between environments that induce the same type of stress (salt vs sorbitol and H_2_O_2_ vs diamide) than between different types of stress. Indeed, we observed a strong correlation of the fitness effects of mutations between salt and sorbitol (r= 0.79) and between H_2_O_2_ and diamide (r= 0.74). However, these correlations are not much different from comparing any other pairs of environments (Supplementary Table 1). Again, diamide is the one environment that stands out with respect to this pattern. Here, the average correlation between diamide and other non-H_2_O_2_ environments is 0.62 (vs r=0.74 between H_2_O_2_ and diamide), suggesting that, in this case, H_2_O_2_ and diamide may share some metabolic features where Hsp90 is involved. One possible explanation for this is that the oxidative stresses of diamide and H_2_O_2_ cause an increase in the expression of chaperones, including Hsp90 (Gasch et al. 2000), inducing similar gene expression responses to those observed during thermal stress in yeast (Gasch et al. 2000).

We also observed a tendency for selection coefficients from the same batch of experiments to be more strongly correlated than selection coefficients inferred from different batches of experiments (Supplementary Table 2). This is not unusual (Venkataram et al 2016) and may be due to small differences in the growth conditions between batches or during the sequencing steps. To mitigate this effect, we performed the analyses to detect beneficial and deleterious hotspots either based on at least two replicates, or using the average of the estimated fitness effects between replicates (as indicated below) and re-categorizing mutations based on the mutant category of both replicates (see also Materials and Methods).

The large correlations observed between most environments suggest that potential costs of adaptation (discussed in detail further below) should be the exception rather than the rule. This is at odds with previous results from a smaller region of Hsp90 (Hietpas et al 2013), where most beneficial mutations detected at high salinity were deleterious in other environments. A potential explanation for this discrepancy lies in the choice of the protein region. Whereas the previous study focused on only nine amino acids and thus chose a set of positions that are likely to show functionally specific patterns, we here had the experimental power to scan a larger region of the protein. Along a larger, less specifically chosen, stretch of the protein, it is likely that many positions and mutations play the same role across many environments. In this case, costs of adaptation of beneficial mutations could still be large, but due to their low proportion in the overall pool of observed mutations they barely affect the correlation of selection coefficients across environments. Indeed, when computing the correlation in a 10 amino acid sliding window, we see high variability in the correlations (Supplementary Figure 7), e.g. with a range between 0.36 and 0.84 for the correlation between Diamide and Standard environments. This suggests that specific regions may represent functionally important positions (see below).

### Beneficial mutations are present across all environments and are enriched in a region of Hsp90 that is implicated in stabilization of interdomains and client binding interfaces

Next, we identified the proportion and identity of putatively beneficial mutations across all treatments (Supplementary Table 1, Supplementary Figure 8, 9). We categorized mutants as putatively beneficial if the lower limit of the 95% confidence interval of the selection coefficient was larger than 0, and we considered as strongest candidates those that overlapped between replicates from the same environment. As reported above, wild-type like mutations are the most abundant category in all environments, with a low proportion but considerable number of beneficial mutations in almost all environments (Supplementary Figure 8, 9). We found the lowest number of beneficial mutations in H_2_O_2_ and salt environments (n_H2O2_=65 and n_Salt_=64, around 2.5% of the mutations) and the highest in diamide (n_Diamide_=307, 12.3%).

The proportion of beneficial mutations, interpreted in the light of Fisher’s Geometric Model (Fisher 1930, Tenaillon 2014), should be informative about the “harshness” of the experimental environments and the resulting potential for adaptation. Specifically, Fisher imagined that populations evolve in a multidimensional geometric phenotype space, where the same mutation is more likely to be beneficial if it happens in an individual that is far from the phenotypic optimum (Fisher 1930, Tenaillon 2014). Thus, we expected that in environments with large doubling times, corresponding to lower absolute fitness of the reference type (Supplementary Figure 1), we should observe a larger number of beneficial mutations. Moreover, we expected the lowest number of beneficial mutations in the standard environment, to which the wild type is well adapted and in which it has the lowest doubling time. However, this expectation was met only in the diamide environment. At high salinity and in sorbitol, the observed proportion of beneficial mutations was lower than in the standard environment. This discordance with expectations from FGM could be due to the model’s very general assumptions which, for example, include that mutations affect all phenotypic dimensions with equal weight (see, e.g., Harmand et al. 2017). Many of these assumptions are likely violated here, because, for example, Hsp90 plays a very different functional role in the osmotic stress response in comparison to diamide induced oxidative stress - probably resulting in different phenotypic distributions of the same mutations in the (anyways rather abstract) phenotype space.

Specifically, it is known that Hsp90 basal function together with its co-chaperone Cdc37 is required for the induction of the osmotic stress response in yeast via activation of the Hog1 kinase in the HOG pathways (Hawle et al. 2007, Yang et al. 2007). However, increased salinity or sorbitol do not cause an increase in Hsp90 mRNA expression in comparison to diamide exposure, which causes an increase in mRNA expression similar to what is observed during heat shock (Gasch et al. 2000). The decoupling of function and expression, and the low number of beneficial mutations suggest that Hsp90s role during osmotic stress may be related to the activation of the general stress response mechanism of the cell (Mager and Siderius 2002). Another possibility is that osmotic stress does not cause heat shock induced protein unfolding (Richter et al.,2010) or diamide induced protein modifications that may compromise protein and cellular function (Gash et al., 2000). Thus, under osmotic stress conditions the cell would not require increased expression of Hsp90. This suggests that mutations may play distinct roles for different conditions, perhaps under osmotic stress activation of clients and under diamide could be activation of clients and/or increase in Hsp90 expression and function.

Interestingly, beneficial mutations in diamide were dispersed across the whole middle domain (Supplementary Figure 8), whereas most of the other environments showed a clear enrichment of beneficial mutations in the region of positions 356-366 (Figure 3, Supplementary Figure 8). This hotspot of beneficial mutations was particularly strong in sorbitol (Supplementary Figure 8). Out of the 31 beneficial mutations that showed a beneficial effect in at least two environments, 18 were located in this hotspot.

The beneficial hotspot contains residues that are part of an allosteric center which mediates distinct conformations in structures with and without client (Czemeres et al. 2017, Blacklock et al 2013,2014). Despite its different structural arrangements, the hydrophobic amino acids in this region are mostly buried from solvent (Figure 3B), indicating that the primary role of this region is to mediate the stability and rearrangement of different conformations. One position that particularly stands out with respect to the number of beneficial mutations is 364. Position 364 lies at the center of the cluster of the 10% largest average proportions of beneficial mutations (Figure 3A) and shows a clear deviation from the usually observed shape of the DFE (Supplementary Figure 10). Specifically, we identified 6 mutations at this position that are beneficial in at least two environments (Supplementary Figure 10). The burial of the large hydrophobic side chain of F364 should provide local stability. The beneficial mutations at position 364 and in this region in general suggest that disruption of local stability and conformational dynamics may alter Hsp90 function. Because this beneficial region partially overlaps with an allosteric center involved in Hsp90 conformational stability and dynamics (Blacklock et al., 2013, 2014) we hypothesize that Hsp90 function may be altered by the identified candidate beneficial mutations in a manner that disrupts local stability and dynamics of this region. Furthermore, these disruptions may promote conformational changes in the neighboring client binding loop that allow this region to sample a larger and/or different conformational space in a manner that changes the relative affinity and thereby priority of different clients - a property that we speculate could provide benefits in specific conditions. Consistent with this hypothesis, the beneficial hotspot is adjacent to a known client binding loop (Figure 3) such that alterations in the structure or dynamics of the hotspot are likely to influence client binding nearby. It is possible that natural selection balances client priorities integrated over multiple conditions, which provides opportunities for Hsp90 mutations to improve priorities for individual conditions (Flynn et al 2020).

### The proportion of deleterious, wild-type like, and beneficial mutations varies greatly along the protein sequence

We showed above that the overall correlations of fitness effects across environments were generally large, which indicates similar effects of the same mutations across environments. In contrast and similar to the local correlations of fitness effects (Supplementary Figure 7), the proportions of beneficial, wild-type like, and deleterious mutations vary greatly along the protein sequence. Whereas the overall pattern is similar between environments, the relative proportions differ between environments (Supplementary Figure 8, 9). Our finding that the same positions are enriched for deleterious and beneficial mutations across environments suggests that the structural properties of the middle domain in these regions, rather than its binding partners, might be the most important factor for predicting the fitness effects of mutations.

Interestingly, the regions with the largest proportions of deleterious mutations are located near the ends of the beneficial hotspot region (position 347 and 376-385, respectively) (Supplementary Figure 8). At the latter deleterious hotspot, around 70% of all mutations are deleterious in a sliding window of 10 amino-acid positions. Such a shift of the DFE towards deleterious mutations suggests that this protein region is under strong purifying selection. Structurally, the region 376-385 is part of a catalytic loop required for ATP hydrolysis which is necessary for the activation of all known clients (Wolmarans et al. 2016). Residue R380 in the catalytic loop binds to and stabilizes the leaving phosphate of ATP (Figure 3C) and mutations at this position compromise Hsp90 function and cell viability (Meyer et al.2003). The efficiency of catalysts depends strongly on geometry, and the precise location of R380 relative to ATP is likely tightly linked to Hsp90 function. The regions adjacent to R380 appear to be important for positioning the catalytic arginine, providing a rationale for the strong purifying selection that we infer. The second deleterious hotspot at position 347 is also located at the interface of the ATPase domain and the middle domain (Figure 3D), consistent with current understanding that Hsp90 mechanism requires precise ATP-dependent interactions between these domains (Schopf et al. 2017).

### Few mutations show costs of adaptation

Among the identified beneficial mutations, we were specifically interested in those that are deleterious in other environments, which results in a so-called “cost of adaptation”. The same phenomenon has also been termed antagonistic pleiotropy. In previous work we reported a large prevalence of costs of adaptation in a 9-amino-acid region of Hsp90 (Hietpas et al 2013). In this study, we found that costs of adaptation were not pervasive (Figure 4, Supplementary Table 3). Consistent with the comparatively low correlations of fitness effects and the special role of the diamide stress discussed above, we observed the largest proportion of mutations that show costs of adaptation between diamide and other environments. Whereas the mean proportion of mutations displaying costs of adaptation across all comparisons of environments was 14.3%, comparisons that included diamide showed on average 22.7% mutations with costs of adaptation.

Mapping the proportion of beneficial mutations that are deleterious in other environments along the protein sequence, we observed a hotspot for costs of adaptation at amino-acid positions 381-391 (Figure 5). Interestingly, this region showed costs of adaptation across various environments, which indicates that each environment has specific beneficial mutations which are deleterious in other environments. Structurally, this region also belongs to the catalytic loop involved in ATP hydrolysis discussed above. Indeed, the identified region partly overlaps with the above-discussed hotspot of deleterious mutations (positions 376-385), which is unsurprising since a larger proportion of deleterious mutations also increases the (statistical) proportion of mutations that can be classified as costly.

We next computed whether there was a correlation between the effects of the subset of beneficial mutations in one environment with their effect in the other environments. Indeed, the effects of beneficial mutations in diamide were negatively correlated with the same mutation’s effect in all other environments (Figure 5, Supplementary Figure 11) except H_2_0_2_. In other words, the more beneficial a mutation was in diamide, the more deleterious it tended to be in salt, standard and sorbitol. Again, this suggests that the beneficial mutations we identified in diamide may be involved in the response to a very specific type of stress. Across all other environments, we did not see any evidence that stronger beneficial mutations tended to have a more deleterious effect in other environments. In fact, there is a suggestive positive correlation between fitness effects of beneficial mutations with their respective effect in all other non-diamide environments, which points to the presence of synergistic pleiotropy. While generally defined as mutations having the same effect on more than one trait, in the context of this study synergistic pleiotropy would mean that beneficial mutations tended to have a similar ranking across environments. This is in line with results from studies of *E. coli* using experimental evolution (Ostrowski et al 2005, Dillon et al 2016) or mutation accumulation (Sane et al 2018). These studies reported synergistic pleiotropy between different carbon source environments, and rare presence of antagonistic pleiotropy. Specifically, Sane et al (2018) found that this pattern was maintained also for the categories of neutral and deleterious mutations, and that mutations with larger effect were more likely to show antagonistic pleiotropy. The reported prevalence of synergistic pleiotropy was associated with a sharing of the metabolism and transport of resources (Ostrowski et al 2005, Dillon et al 2016, Sane et al 2018).

## Conclusion

Recent advances in experimental and technological approaches have led to the feasibility of large-scale screens of the DFE of new mutations, which, from an evolutionary point of view, is a key entity to determine the potential for adaptation. Here, we took a new route by mapping the proportions of beneficial, wild-type like, and deleterious mutations along a 119-amino-acid region of the Hsp90 protein in yeast, a protein that is heavily involved in the response to environmental stressors. This approach allowed us to create a genotype-phenotype map to identify specific protein regions that may be important for adaptation to new environments. Specifically, by comparing the DFE along the protein sequence and between environments, we identified hotspots of beneficial and deleterious mutations that are shared between environments, and a region in which beneficial mutations in one environment tend to be deleterious in other environments. Interestingly, neither of these regions coincided with the best described client binding loop in the studied region, which we had *a priori* considered the most likely candidate to display patterns different from the rest of the region. Moreover, our analyses suggest that mutational effects generally differ little across environments except in diamide, which stood out both with respect to the number and also the distribution of beneficial mutations along the studied protein region. Altogether, our study of the DFE across environments sheds light on the evolutionary role of a specific protein region from a new perspective.

## Materials and Methods

### Generating point mutations

To accurately measure the fitness effects of all possible point mutations in a large portion of the middle (client-binding) domain of yeast Hsp90, we used saturation mutagenesis at positions 291-409. We used a cassette ligation strategy to introduce mutations as previously described (Hietpas et al. 2012). This strategy reduces the likelihood of secondary mutations because it avoids the potential for errors during PCR steps. As a control for the mutational procedure, twelve positions were randomized in isolation and Sanger sequenced to assess the level of incorporation of all four nucleotides at each position in the target codon. At all randomized nucleotide positions within these twelve samples, we observed a similar magnitude of signal for each of the four nucleotides. For larger scale production, libraries were generated at 10 positions at a time as previously described (Hietpas et al. 2012). As additional controls, we generated a sample containing individual stop codons as well as the parental wild-type Hsp90 sequence. All variants of Hsp90 were generated in a plasmid (pRS414GPD) previously shown to produce endogenous levels of Hsp90 protein in yeast (Chang and Lindquist 1994).

### Constructing barcoded libraries

To improve the efficiency and accuracy of fitness estimates, we added barcodes to a non-functional region of the plasmid, ∼200 NTs downstream from the 3’ untranslated region of Hsp90. Barcodes were introduced using a cassette ligation strategy (Supplementary Figure 12). Plasmid libraries were treated with NotI and AscI to generate directional sticky ends. NotI and AscI recognize 8-base cut sites that are unlikely to cut any of the Hsp90 variants in the library. We designed and annealed barcode forward and barcode reverse oligos together such that the resulting duplex product included a central N18 region bracketed by constant regions that facilitate annealing and overhangs that direct directional ligation into NotI and AscI overhangs. One of the constant ends in the designed oligo cassette contained an annealing region for an Illumina sequencing primer. Barcoded libraries were transformed into *E. coli* and pooled into a bulk culture that contained about 10-fold more transformants than Hsp90 variants in the library. We purified barcoded plasmids from this bulk culture. This procedure resulted in approximately 20 barcodes for each Hsp90 codon variant in the library (Supplementary Figure 13). The potential diversity in the N18 barcode that we used (4^18^ ∼ 10^11^) far exceeds the number of barcodes that we utilized (∼64*119*10∼10^5^), which makes it likely that each Hsp90 variant receives a barcode that differs from all other barcodes at multiple nucleotides. With this setup, errors in sequencing of barcodes can be detected and eliminated from further analysis, which reduces the impact of sequencing misreads on estimates of variant frequency and fitness. Additional controls consisting of individual stop codons and wildtype Hsp90 were barcoded separately. For these controls, we isolated barcoded plasmid DNA from individual bacterial colonies and determined the barcodes by Sanger sequencing (Supplementary Figure 12).

### Associating barcodes to mutants

To identify the barcodes associated with each Hsp90 variant in our libraries, we used a paired-end sequencing approach essentially as previously described (Hiatt et al. 2011, see also Supplementary Figure 14). Using paired-end sequencing on an Illumina MiSeq Instrument barcodes were associated with variant genotypes via reading from the Hsp90 gene with a 250 base-pair read and the associated N18 barcode with a 50 base-pair read. To reduce the size of the DNA fragments for efficient Illumina sequencing, we removed a portion of the Hsp90 gene such that the randomized regions were closer to the N18 barcode. This was done to increase the density of DNA on the sequencer by reducing the radius of clonal clusters during sequencing. Plasmid DNA was linearized with StuI and NotI endonucleases that removed ∼400 bp’s of DNA. The linearized products were circularized by blunt ending followed by ligation at DNA concentrations that predominantly lead to unimolecular ligations (Revie et al 1998). The resulting DNA products were amplified using a PE2_F primer and the standard Illumina PE1 primer that anneals next to the N18 barcode. Two PE2_F primers were designed in order to read across the region of Hsp90 that we randomized. PCR products were gel purified and submitted for paired end sequencing. We obtained sufficient paired-end reads such that the average barcode was read more than 10 times (Supplementary figure 13). The paired-end sequencing data was subjected to a custom data analysis pipeline to associate Hsp90 variants with barcodes. First, very low-quality reads with any Phred score less than 10 in reads 1 or 2 were discarded. Next, the data were organized by the barcode sequence. For barcodes with three or more reads, we constructed a consensus of the Hsp90 sequence read. We compared the consensus sequence to the parental Hsp90 sequence in order to determine mutations. Consensus sequences containing more than one protein mutation were discarded. The pipeline output generated a file organized by point mutation that lists all barcodes associated with that mutation (Supplementary Data). This file was used as the basis for calling a variant based on barcode reads.

### Yeast competitions

As in previous work (Hietpas et al. 2012, Hietpas et al. 2013), we performed Hsp90 competitions using a shutoff strain of *S. cerevisiae* (DBY288) (Supplementary Figure 14). The sole copy of Hsp90 in DBY288 is driven by a galactose-inducible promoter, such that the strain requires galactose for viability and cannot grow on glucose. The introduction of a functional Hsp90 variant driven by a constitutive promoter rescues the growth of DBY288 in glucose media. We introduced the library of middle domain variants of Hsp90 driven by a constitutive promoter into DBY288 cells using the lithium acetate method (Gietz and Schiestl 2007). The efficiency of transformation was more than 10-fold higher than the number of barcodes, such that the average barcode was transformed into more than 10 individual cells. To enable the analyses of full biological replicates, transformations were performed in replicate such that a separate population of yeast were transformed for each biological replicate. Transformed yeast were initially selected in RGal-W (synthetic media that lacked tryptophan and contained 1% raffinose and 1% galactose supplemented with 100 µg/mL ampicillin to hinder bacterial contamination) to select for the plasmid, but not function of the plasmid encoded Hsp90 variant. This enabled us to generate a yeast library containing Hsp90 variants that could support a full range of fitness from null to adaptive. Cells were grown for 48 hours at 30°C in liquid SRGal-W until the culture was visibly opaque compared to a control transformation sample that lacked plasmid DNA but was otherwise identical to the library sample.

To initiate the competition, cells were transferred to shut-off conditions and different environmental stresses. In order to deplete the pool of wild type Hsp90 protein, the library sample was diluted into SD-W (synthetic media lacking tryptophan with 2% glucose and 100 µg/mL of ampicillin) and grown at 30 °C for 10 hours. After depletion of wild type Hsp90 protein, the cells were split into different stress conditions in SD-W media at 30 °C: salt stress (0.5M NaCl), osmotic stress (0.6M sorbitol), oxidative stress (0.6 mM hydrogen peroxide or 0.85 mM diamide), as well as a control non-stress condition. We used a custom built turbidostat (Supplementary Figure 14) to provide constant growth conditions during this competition phase of the experiment. Cells were grown under consistent density and population size. 10^9^cells were maintained in a 50 mL volume over the course of 40 hours for each condition. Rapid magnetic stirring was used to provide aeration to the media and cells. Samples of 10^8^cells were collected after 0, 4, 8, 12, 24, 32, 40, and 48 hours of competition in the different conditions. At the time of collection, samples were centrifuged and the pellets immediately frozen at -80 °C.

### Sequencing of competition samples

The fitness effects of Hsp90 variants were estimated based on frequencies observed in next-generation sequencing analyses essentially as previously described (Hietpas et al. 2012). DNA was isolated from each timepoint sample as described (Hietpas et al. 2013) and the barcodes were amplified and sequenced. Barcodes were amplified using the standard Illumina PE1 primer that anneals next to the N18 barcode and custom designed barcode_forward primers. A set of barcode_forward primers were designed with identifier sequences that could be read during sequencing and used to distinguish each timepoint sample (Supplementary Figure 14C). Each identifier sequence was eight nucleotides in length and differed by at least two bases from all other identifier sequences. Twenty cycles of PCR were sufficient to generate clear products from all samples. These PCR products were purified on silica columns (Zymo Research) and sequenced for 100 base single reads on an Illumina NextSeq instrument. The barcode corresponded to the first 18 bases and the identifier at positions 91-98. The resulting fastq files were processed and analyzed using customized software tools. First, poor quality reads containing any positions with a Phred score less than 20 were discarded. Reads were tabulated if the barcode matched a barcode associated with a point mutation and if the identifier matched with a timepoint. The analyses scripts output a file with the number of reads for each amino acid point mutant at each timepoint in each condition (Supplementary Data). Mutants with 50 or less total reads at the first time point were removed from the data set.

### Estimation of selection coefficients

Inference of selection coefficients was performed via log-linear regression as described in Matuszewski et al 2016 (see also Supplementary Material). To improve estimation accuracy by incorporating information from each individual barcode, the linear model for each amino-acid changing mutation included barcode identities as nominal variables. For each amino-acid changing mutation we obtained a selection coefficient and 95% confidence interval (CI) of the estimate, representing the variation within amino acid due to differences in codon*Barcode tag throughout time. This reduces the impact of potential outliers and averages over synonymous codons within amino acid. Finally, we normalized all selection coefficients by subtracting the median of all mutations that were synonymous to the wild type. This ensures that the average of the selection coefficients of wild type synonyms represents a selection coefficient of 0. To categorize mutations as beneficial, deleterious or wild-type we tested the overlap of the CI with 0. Namely, if the lower CI limit was larger than 0, a mutation was considered beneficial, and if the upper CI limit was smaller than 0, a mutation was considered deleterious. All other mutations were considered wild-type like. We use this terminology to distinguish between neutral and wild-type like mutations. Specifically, we expect that at large population sizes the number of wild-type like mutations is larger than the number of neutral mutations. This is because neutral mutations should behave neutrally with respect to the neutral or nearly-neutral theories, i.e., when *2Ns*<1, where *s* denotes the selection coefficient and *N* the population size (Ohta 1992). Considering the large experimental population sizes of yeast, this threshold for neutrality is far below the measurement accuracy of our experimental setup. Therefore, we expect that neutral mutations are a subset of the wild-type like mutations. Thus, we instead consider as “wild-type like” those mutations that are indistinguishable from the wild type according to our inference accuracy.

### Natural variants

We extracted observed amino acid mutations from Hsp90 sequences of 261 eukaryotic organisms (Starr et al., 2018). Then, for each environment, we compared the distribution of fitness effects of the identified natural variants with the DFE from all mutations. Finally, we also computed the overall proportion of beneficial, deleterious and wild-type like mutations across environments and for the natural variants.

### Costs of adaptation

To compute the costs of adaptation for Figure 4 and 5, we first selected all mutations in a focal environment that were categorized as beneficial. We then computed the proportion of this subset of mutations that was deleterious either in one other environment (for Figure 4) or across all other environments (excluding replicates of the focal environment for Figure 5). Calculating the proportion of mutations with costs of adaptation compensates for variable numbers of beneficial mutations across environments, which may occur due to both biological and experimental reasons (e.g., different error margins). For Figure 2 and 4, all available data sets were considered for the analysis. For Figure 3 and 5, only two replicates of each environment were considered. That was done in order to avoid biases due to the under-or overrepresentation of environments in the analysis.

Analyses were performed using R version 3.5.1 (R Core Team 2018), Mathematica 12.0.0.0, Python 2.7.12 and Pymol 0.99. Code is available as Supplementary Material.

All data will be made available in a data repository upon acceptance of the manuscript.

## Acknowledgements

This work was supported in part by grant R01GM112844 from the National Institutes of Health (to DNAB). IF was supported by a postdoctoral fellowship from the FCT (Fundação para a Ciência e a Tecnologia) within the project JPIAMR/0001/2016. CB is grateful for support by EMBO Installation Grant IG4152 and by ERC Starting Grant 804569 - FIT2GO.

