## Supplementary Material for "The adaptive potential of the middle domain of yeast Hsp90"

#### Supplementary Figures

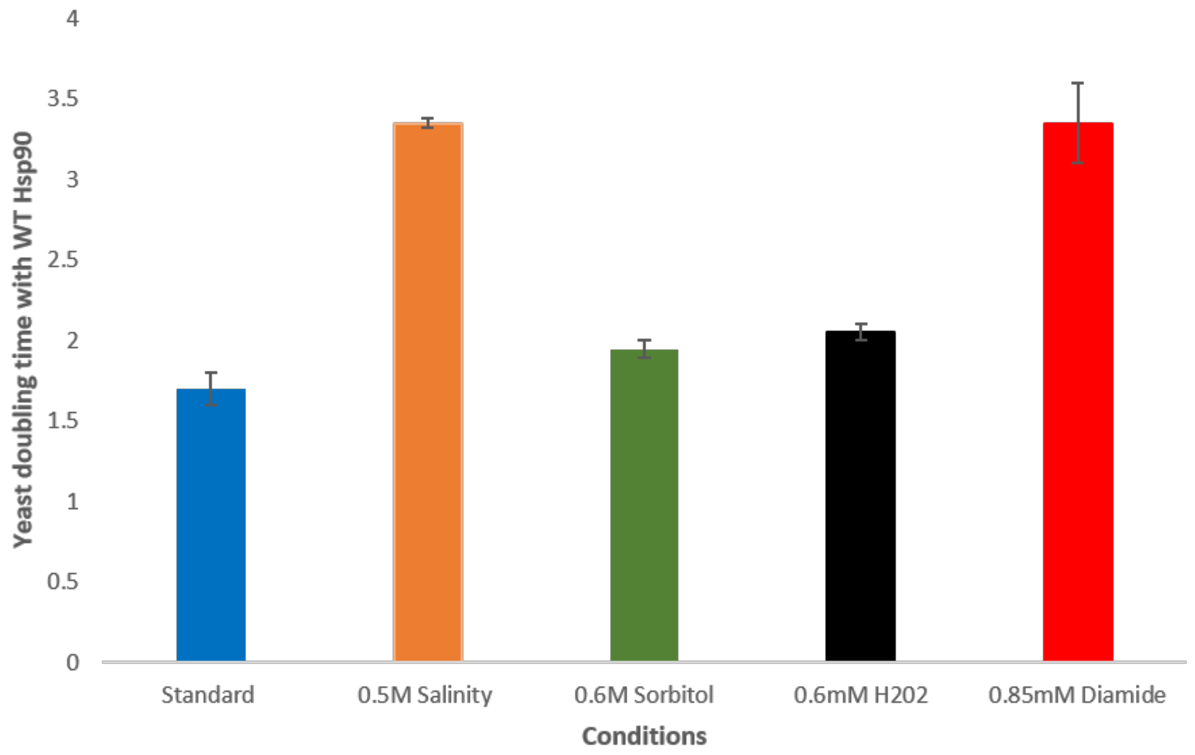


**Supplementary figure 1. Doubling times for WT Hsp90 yeast growth across environments.** In general there is an increase in the doubling time in all environments tested compared to the standard environment. This increase is larger in high salinity and diamide environments, suggesting that these environments exert more stress on the organism.

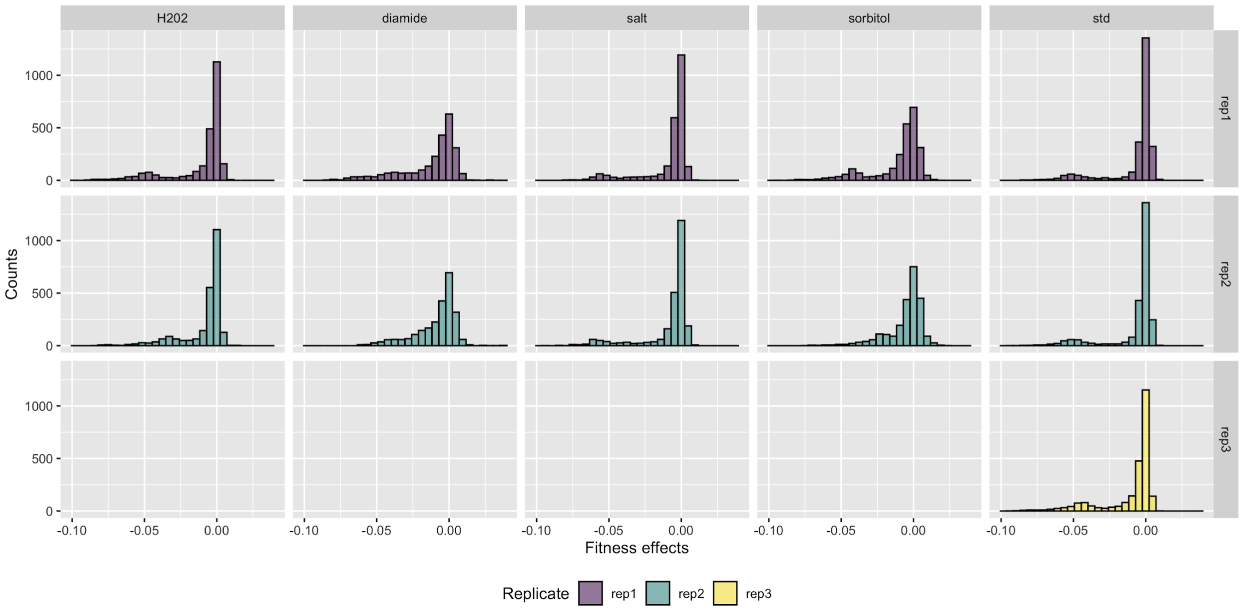


**Supplementary Figure 2 Distribution of Fitness Effects (DFE) across environments.** Replicates within an environment largely overlap. There is a marked difference of the DFE in diamide compared to the other environments. Interestingly, the shape of the DFEs in $H_20_2$and salt is similar to the DFE in the standard environment, with a large number of mutations in the wild-type like region and a smaller peak of deleterious mutations, at which stop codons are located. In diamide and sorbitol, the DFE shows a lower number of wild-type like mutants, a larger variance, and a larger proportion of weakly and strongly deleterious mutations.

### Diamide Replicate 1

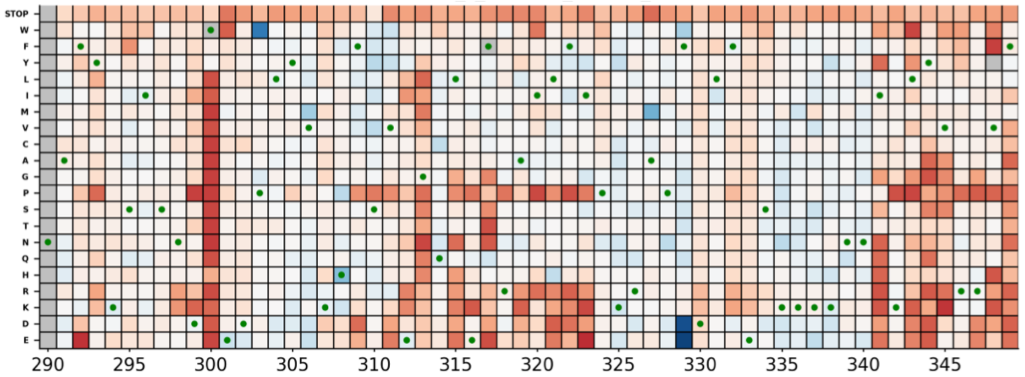


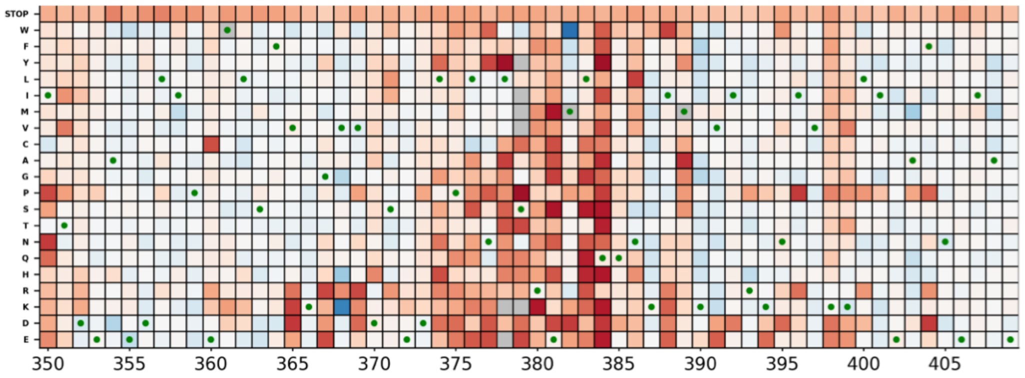


### Diamide Replicate 2

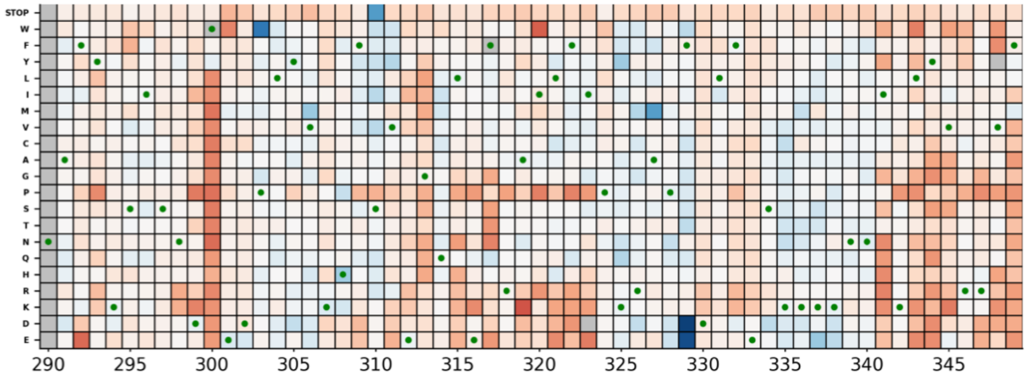


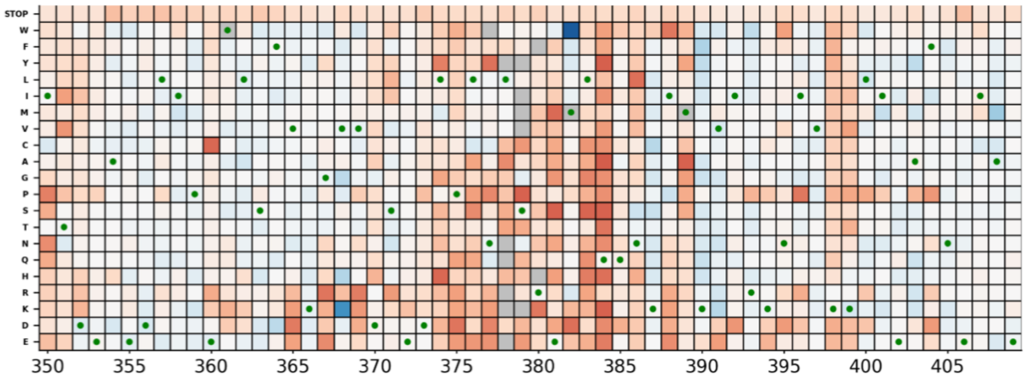

### $H_2O_2$ Replicate 1

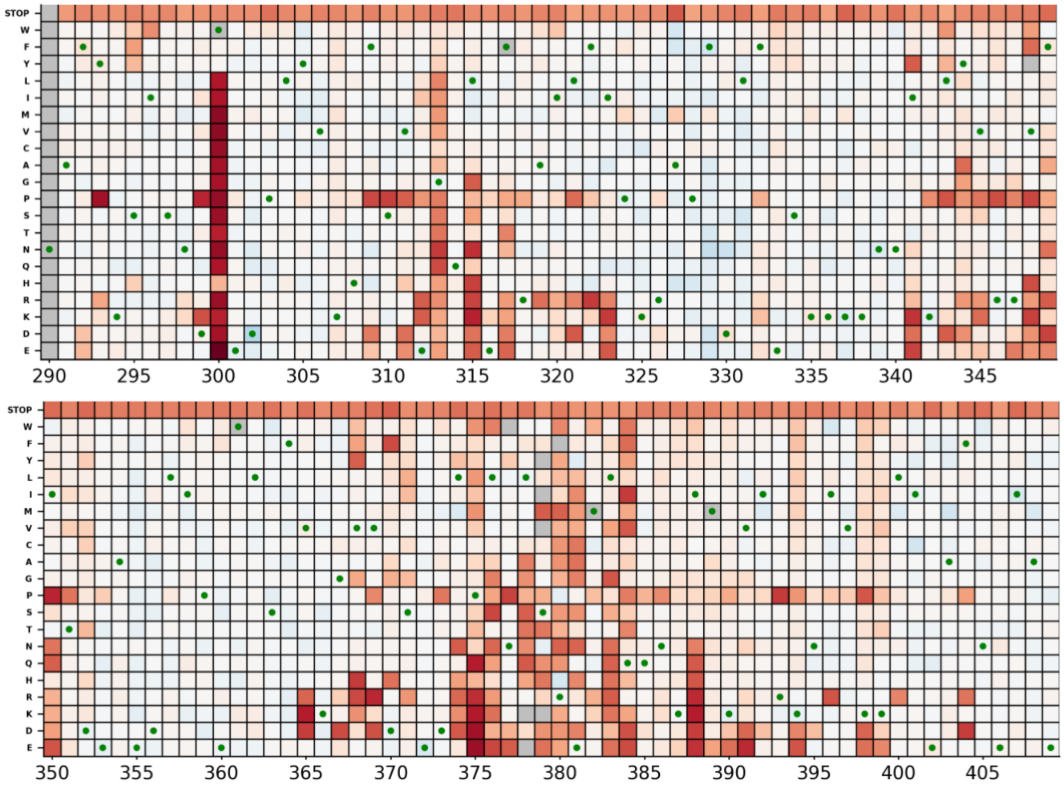


### $H_2O_2$ Replicate 2

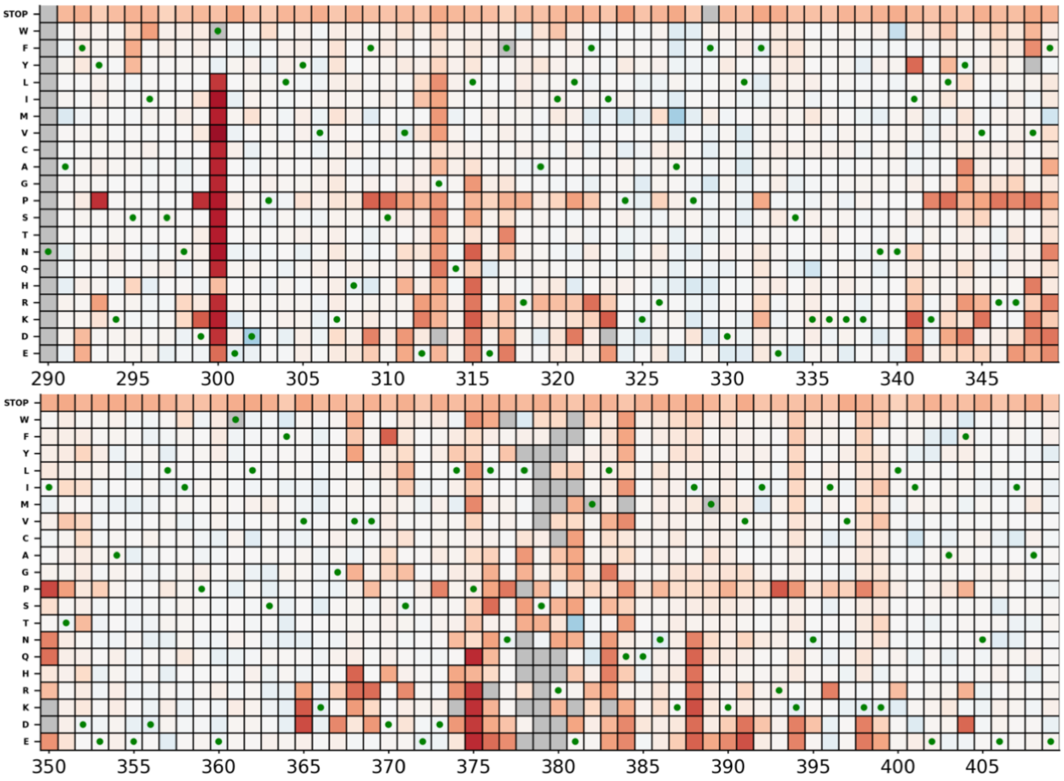

#### High Salinity Replicate 1

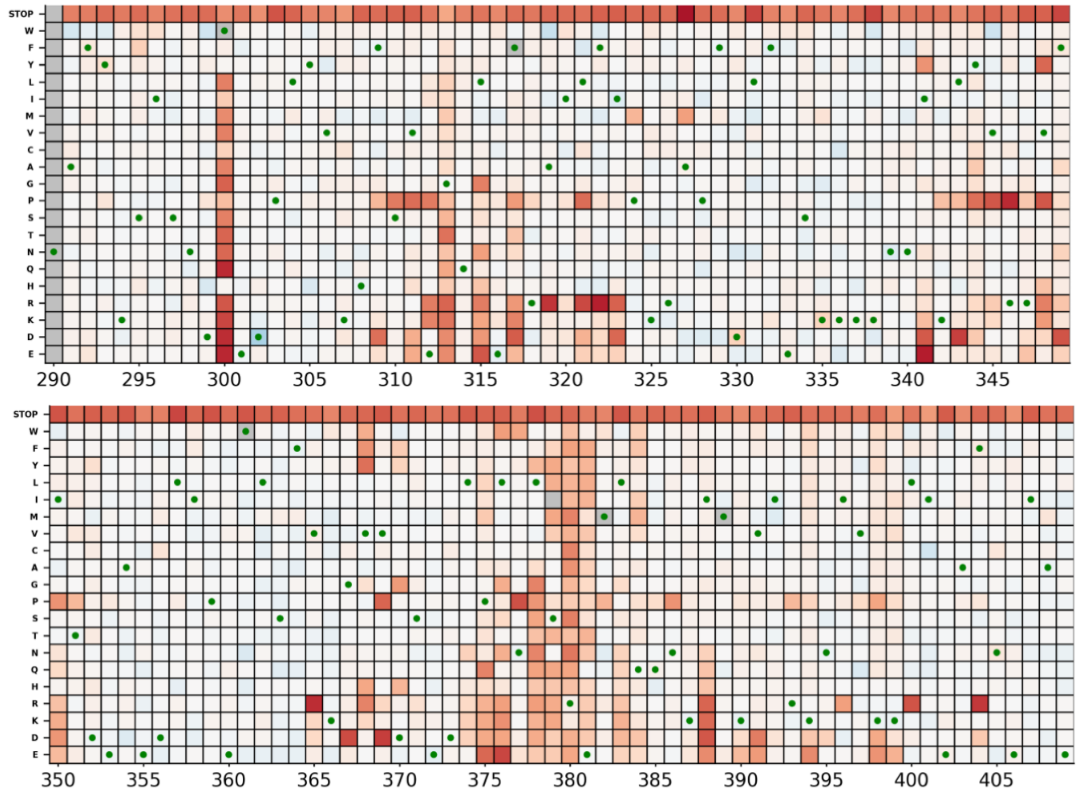


#### High Salinity Replicate 2

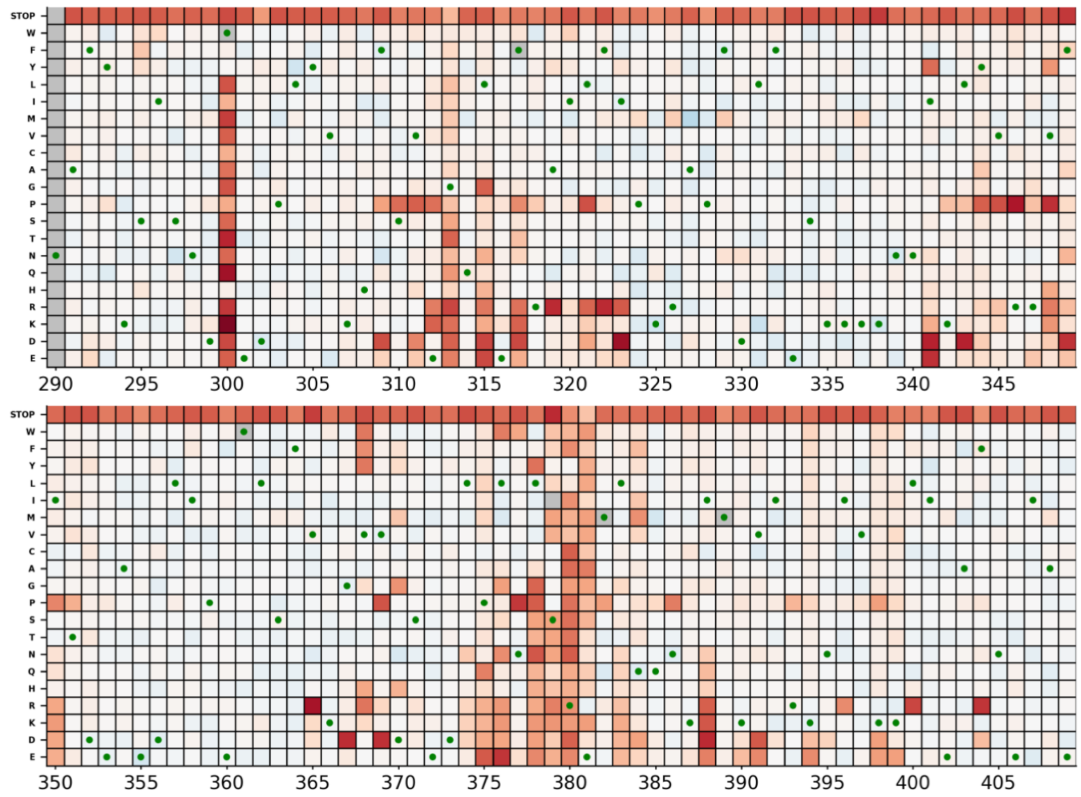

#### Sorbitol Replicate 1

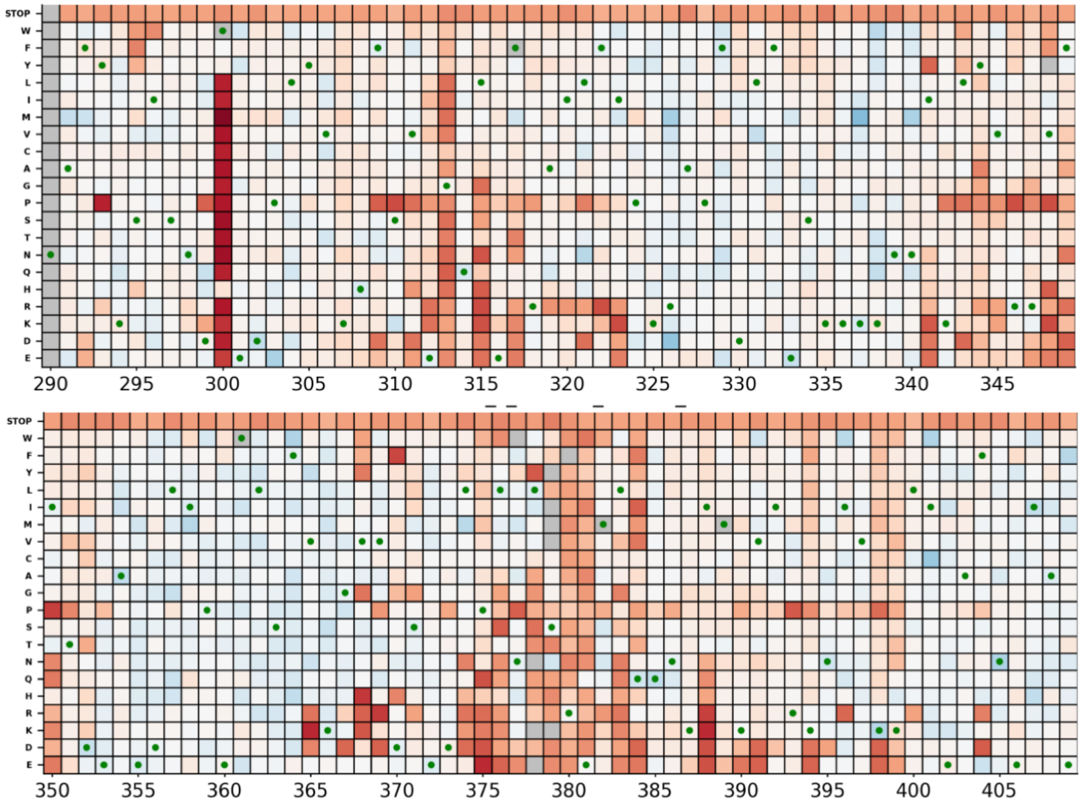


#### Sorbitol Replicate 2

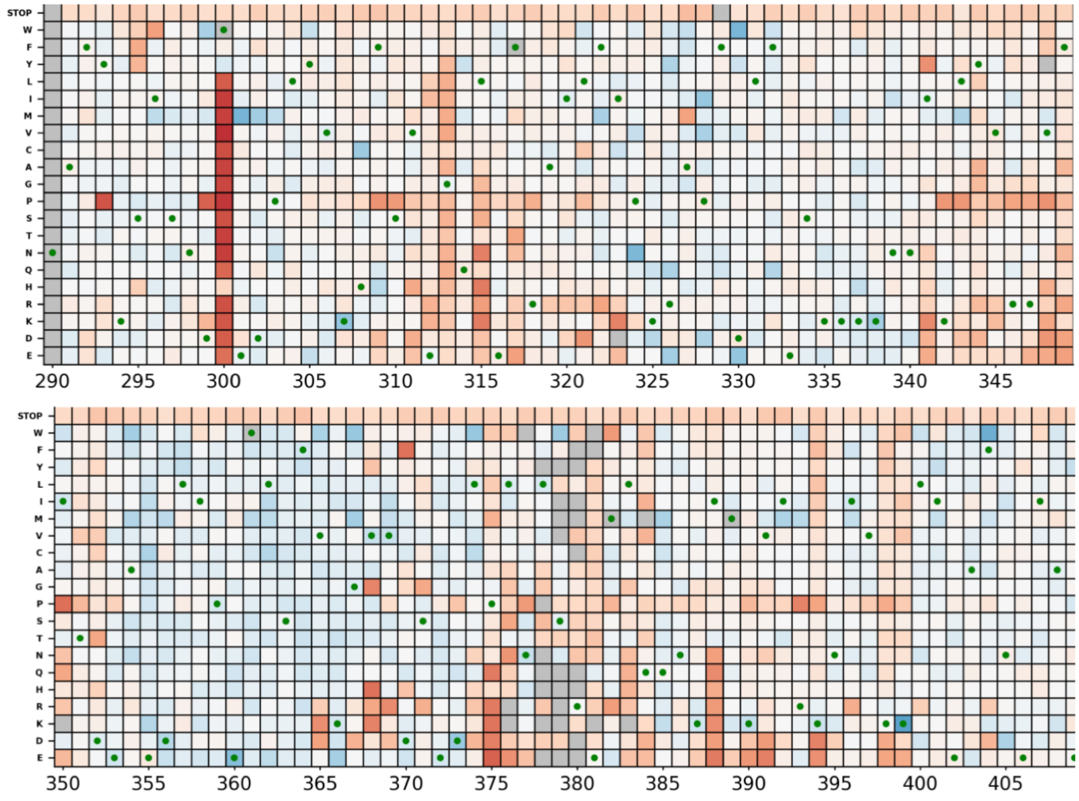

Standard Replicate 1

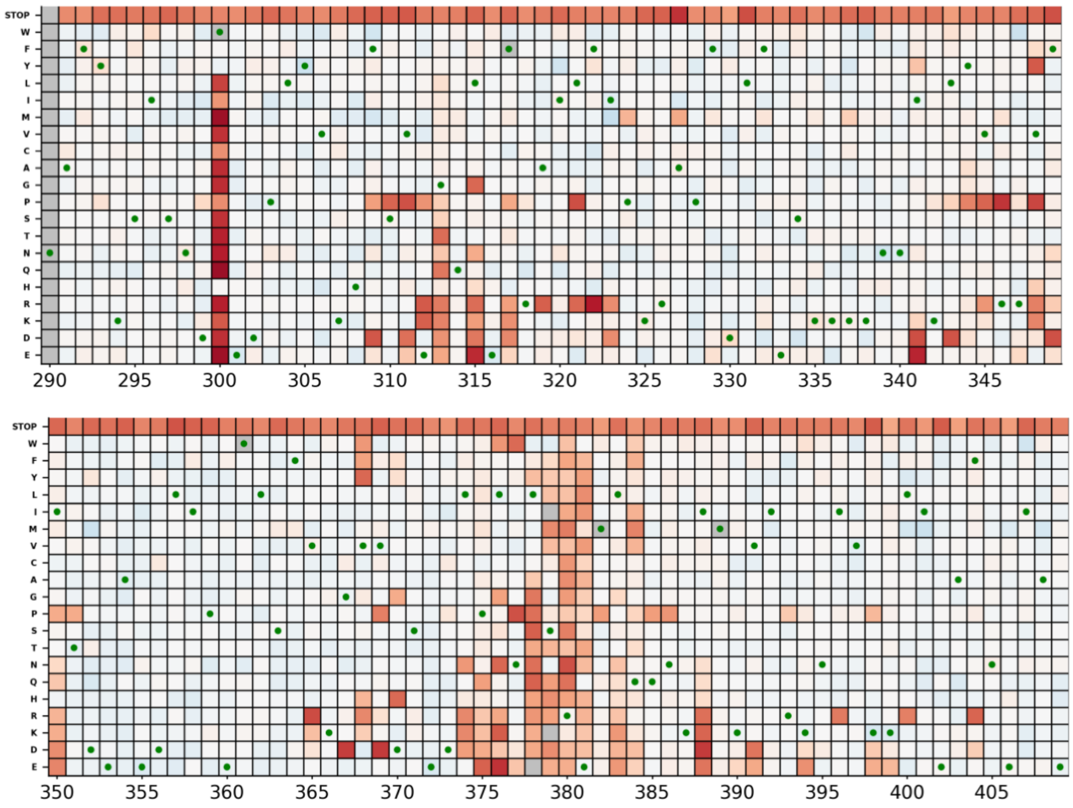

Standard Replicate 2

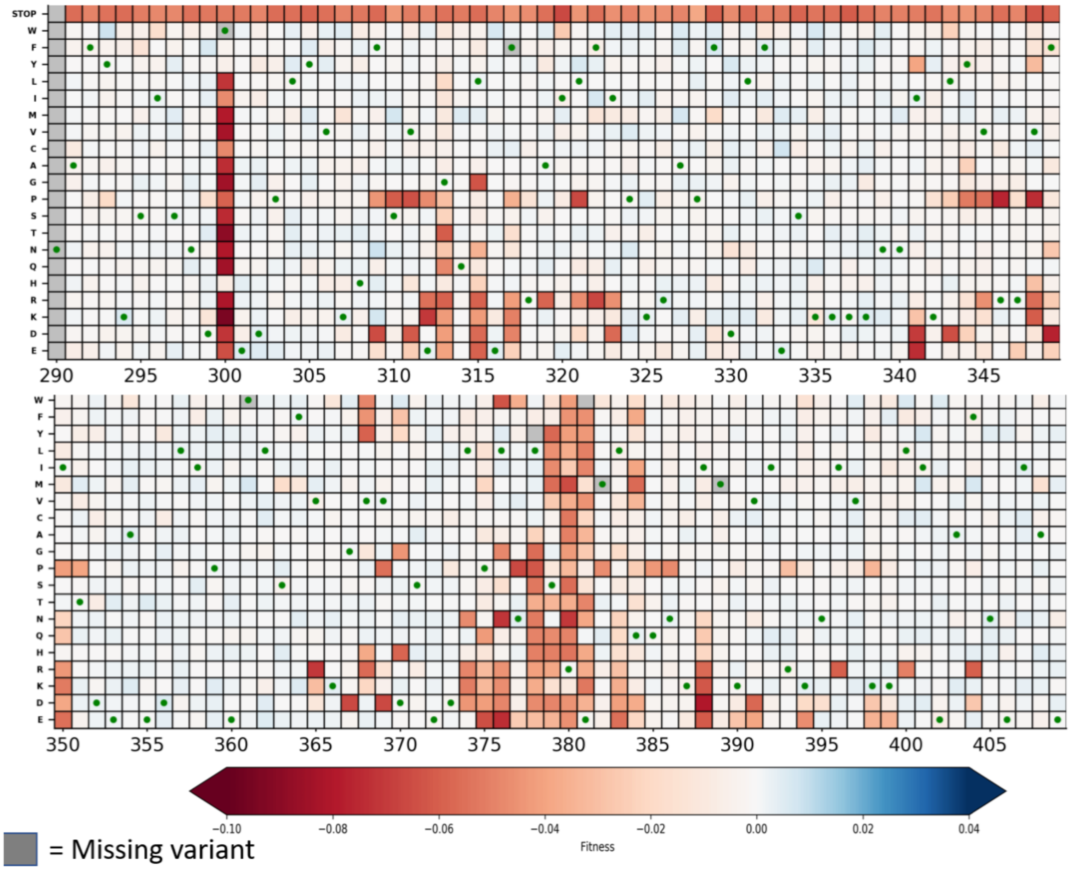

**Supplementary Figure 3. Heatmaps of mutant fitness effects show that the majority of mutations are wild-type like under all conditions, with the exception of amino acid regions ~300, 315-320, 347 and 375-390 that show mostly deleterious mutant fitness effects. Beneficial mutations were found under all conditions but in different regions of the middle domain.** Heatmaps for (A) diamide, (B) $H_20_2$, (C) salt, (D) sorbitol and (E) standard environments.

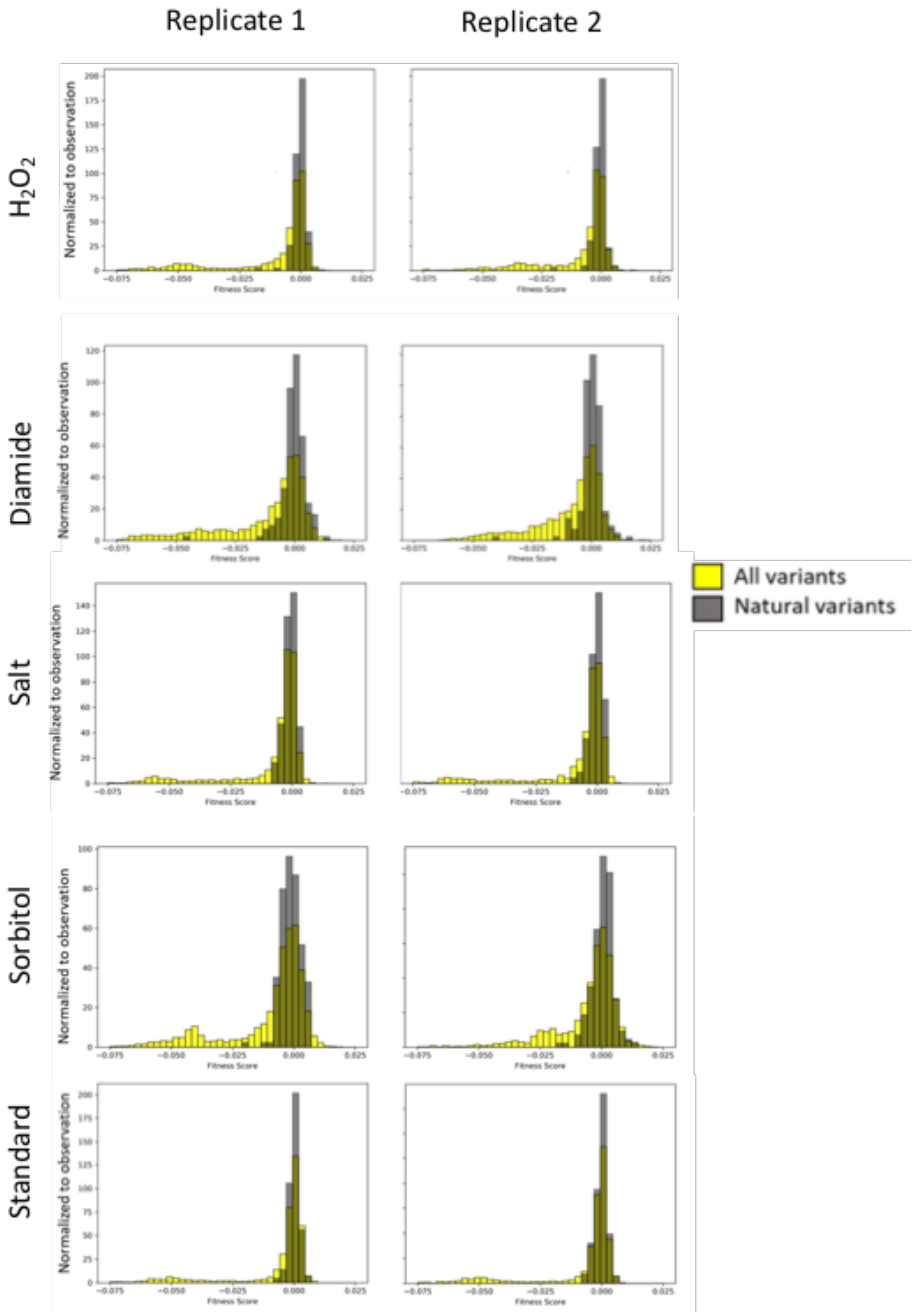


**Supplementary Figure 4**

**Distribution of selection coefficients of mutations found in natural environments vs all mutations studied here.** The x-axis corresponds to the selection coefficient of the mutations and the y axis to counts of mutations normalized to the number of observations. The rows indicate different environments and the columns correspond to 2 replicates for each environment.

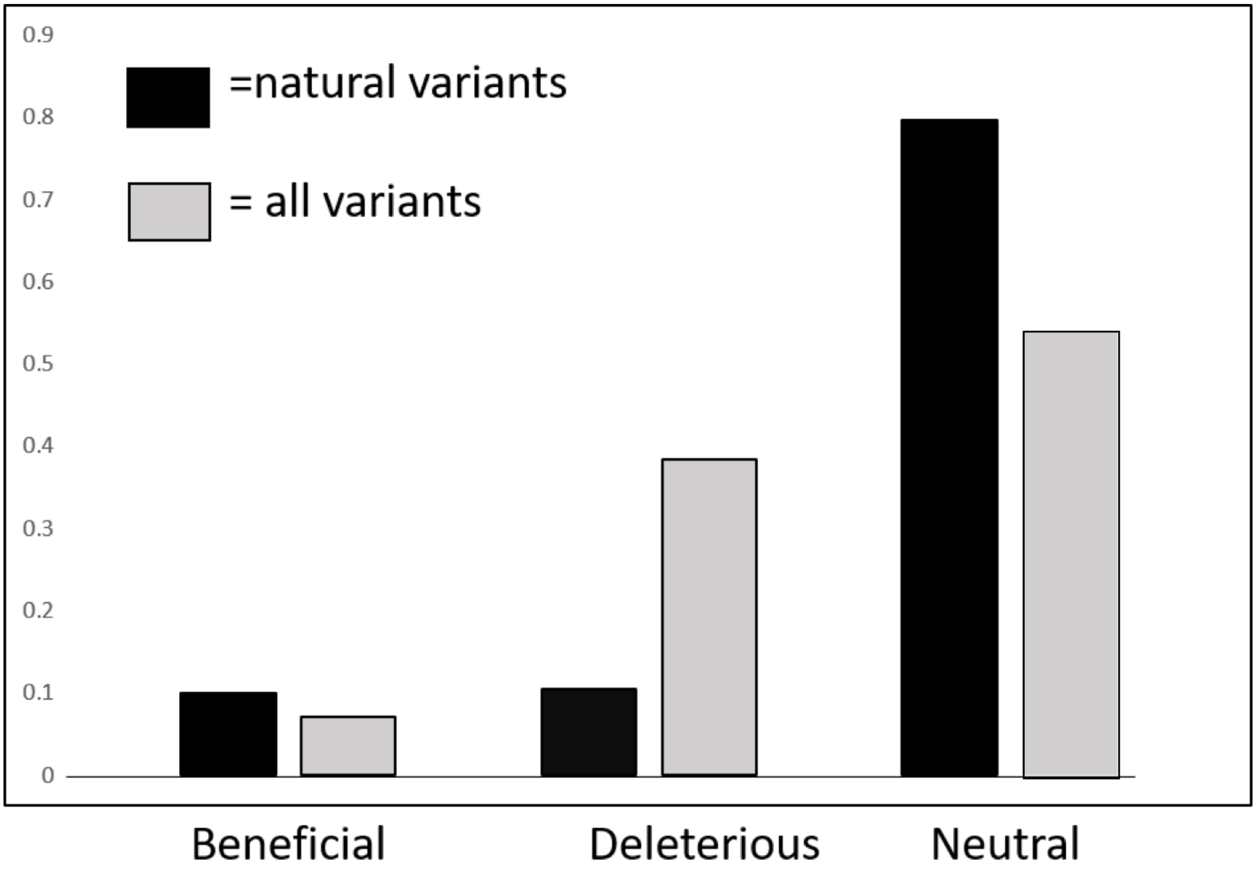


**Supplementary Figure 5**

**Proportion of mutations that are beneficial, deleterious or neutral comparing all mutation our study with the subset that was observed in natural populations of eucaryotes.** The x-axis corresponds to the 3 mutational categories and the y axis to the proportion of mutations in each category.

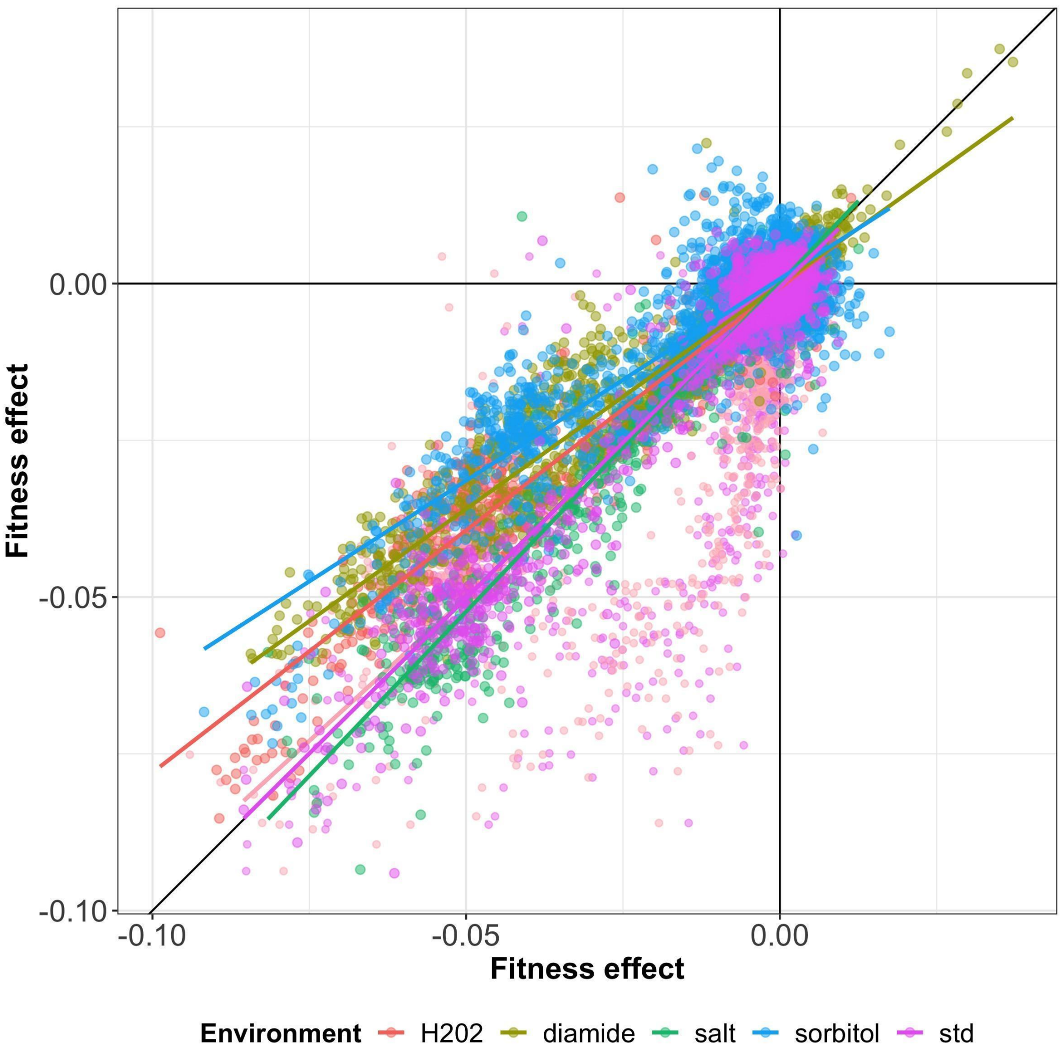


**Supplementary Figure 6**

**Correlation between replicates for each environment.** The x-axis corresponds to the fitness effect of replicate 1 (or replicate 3 for the comparison between standard environment replicate 2 vs 3) and the y axis to the fitness effect of replicate 2 (or 3 for the comparison between standard environment replicate 1 vs 3). In general we observe a very high correlation between replicates within each environment.

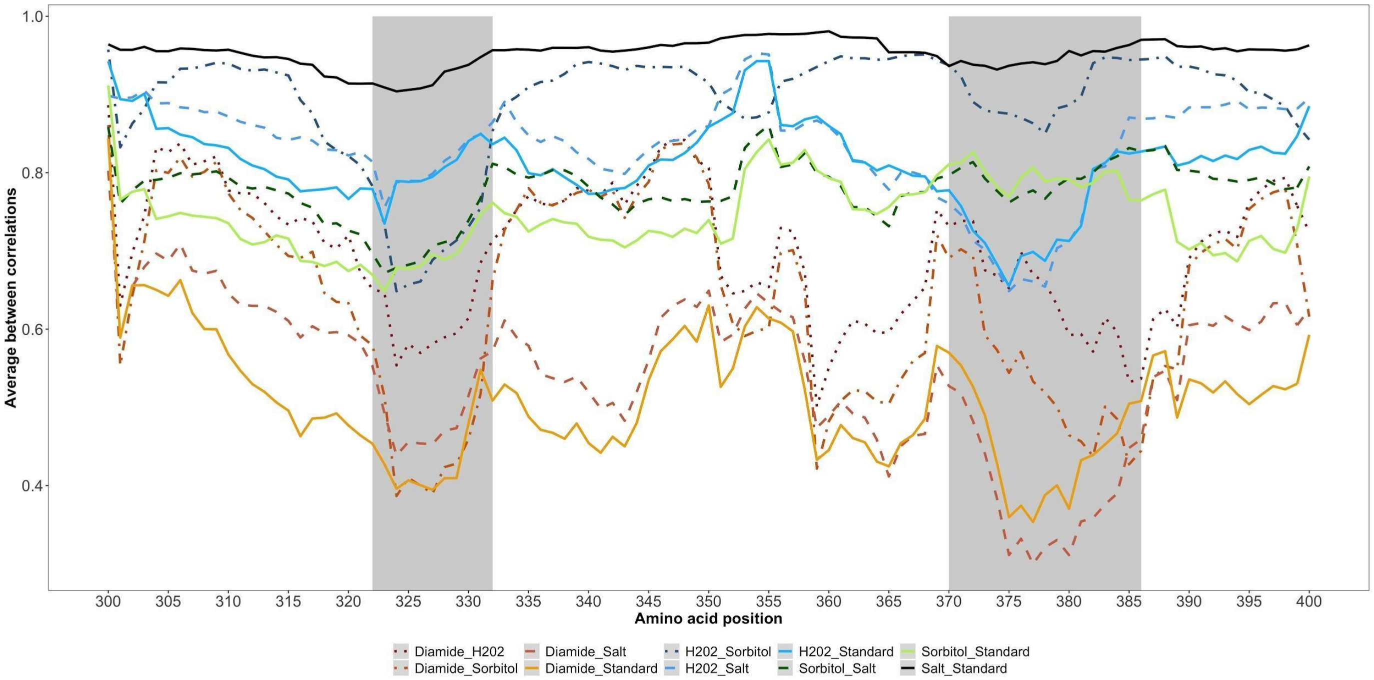


**Supplementary Figure 7**

**Correlation between environments in a 10 amino acid sliding window.** The x-axis corresponds to the amino acid position of the center of the sliding window and the y-axis to the average correlation between environments. Regions with a consistently lower correlation as compared with the baseline for all combinations of environments are highlighted in grey.

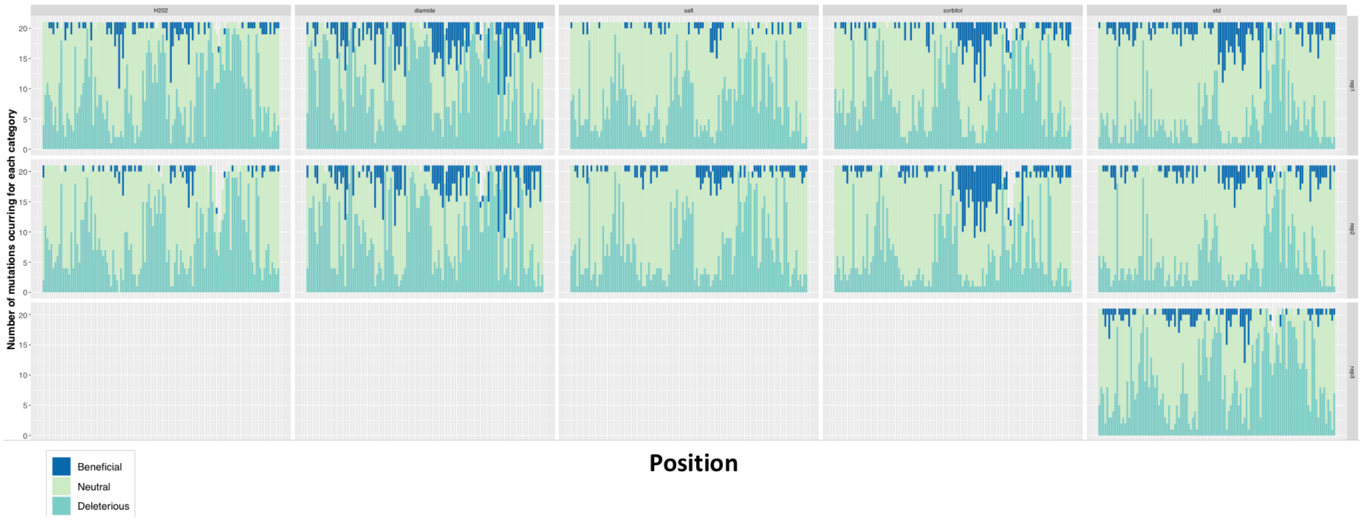


**Supplementary Figure 8**

**Proportions of deleterious, beneficial and wild-type like mutations along the middle domain of Hsp90.** The x-axis corresponds to the amino acid positions studied and the y axis to the proportion of deleterious, beneficial and neutral mutations at each amino acid position. The rows indicate expression level and replicate, the columns the different environments.

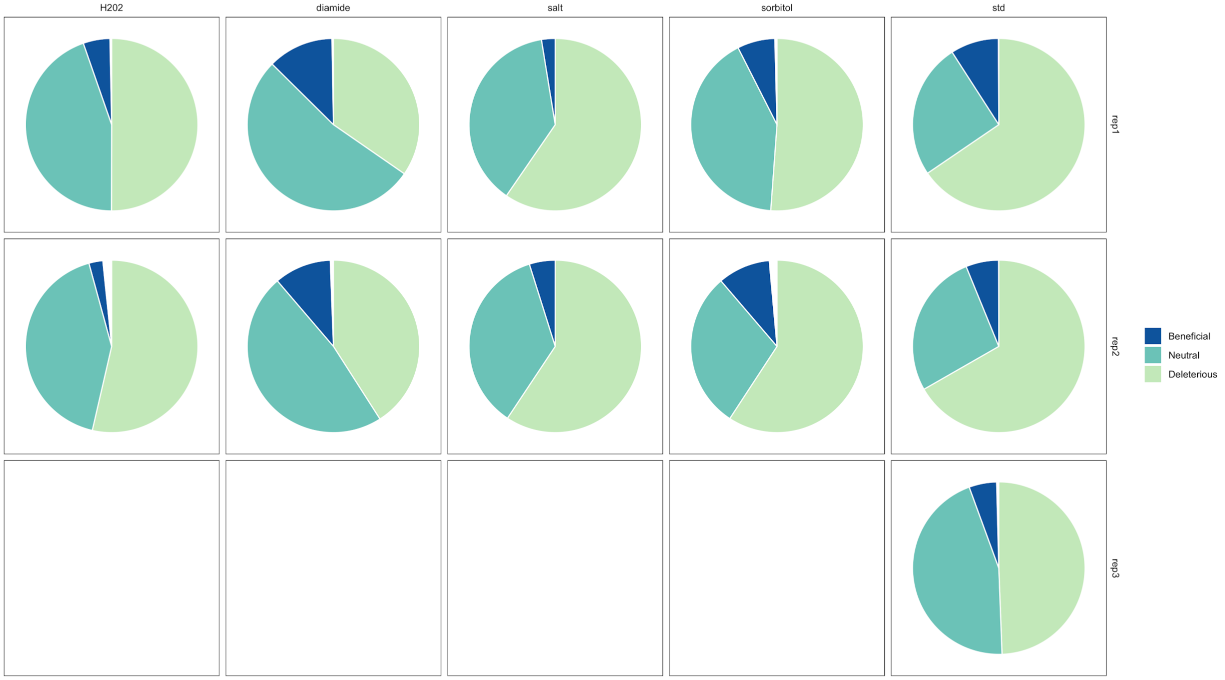


**Supplementary Figure 9**

**Summary of the proportions of beneficial, deleterious and neutral mutations in each environment/replicate**. In general, we see few beneficial mutations, except in diamide. In diamide, there is also a larger proportion of deleterious mutations in comparison with other environments.

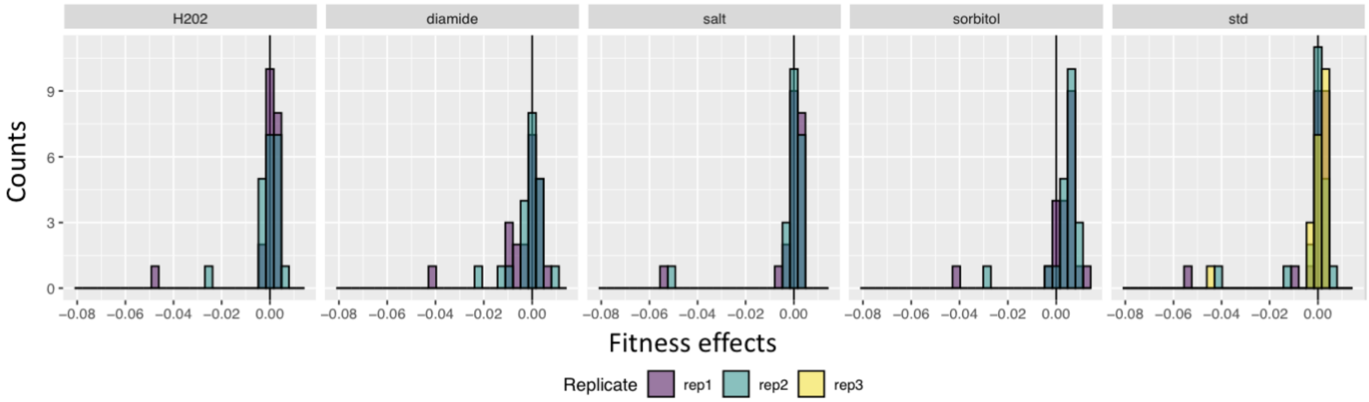


**Supplementary Figure 10**

**Distribution of fitness effects at position 364**. Compared with the DFE of the whole middle domain (Supplementary Figure 2), there is a clear enrichment of beneficial mutations at position 364.

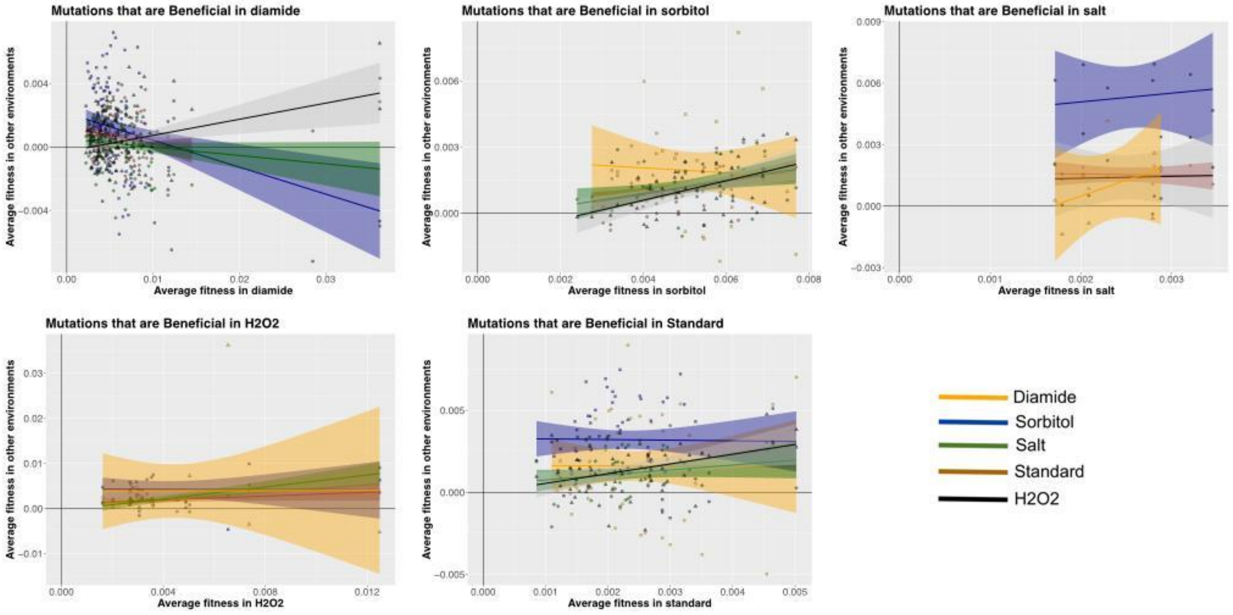


**Supplementary Figure 11**

**Cost of adaptation in different environments.** Each plot represents the average fitness effect of mutations that are beneficial in a specific environment (x-axis) and the average fitness effect in other environments (y-axis). Focal environments are indicated by the color of the lines and points. Note that the x and y scales vary between panels. To improve visualization, we calculated the average fitness effects between the two (or three replicates) and mutations were recategorized based on whether mutations were beneficial in two replicates (beneficial mutations), beneficial in one replicate and neutral in another (beneficial-neutral), neutral in both replicates (neutral mutations), neutral in one replicate and deleterious in the other (neutral-deleterious mutations), deleterious in both replicates (deleterious) or deleterious in one replicate in beneficial in the other (beneficial-deleterious mutations).

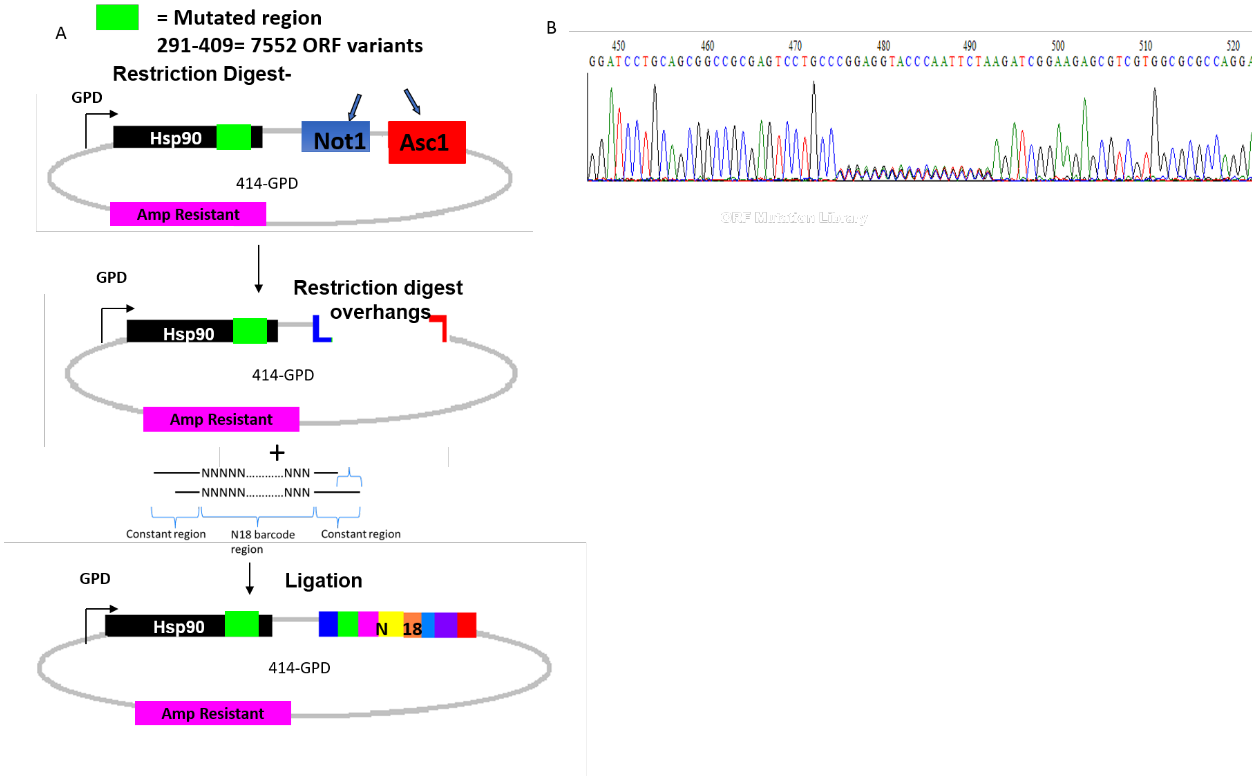


**Supplementary figure 12**. Barcoding mutant library strategy. (A) Schematic of how to incorporate barcodes into plasmid libraries using restriction digests and ligation. (B) Sanger sequence TRACE result of barcoded region of the library at positions 475-493.

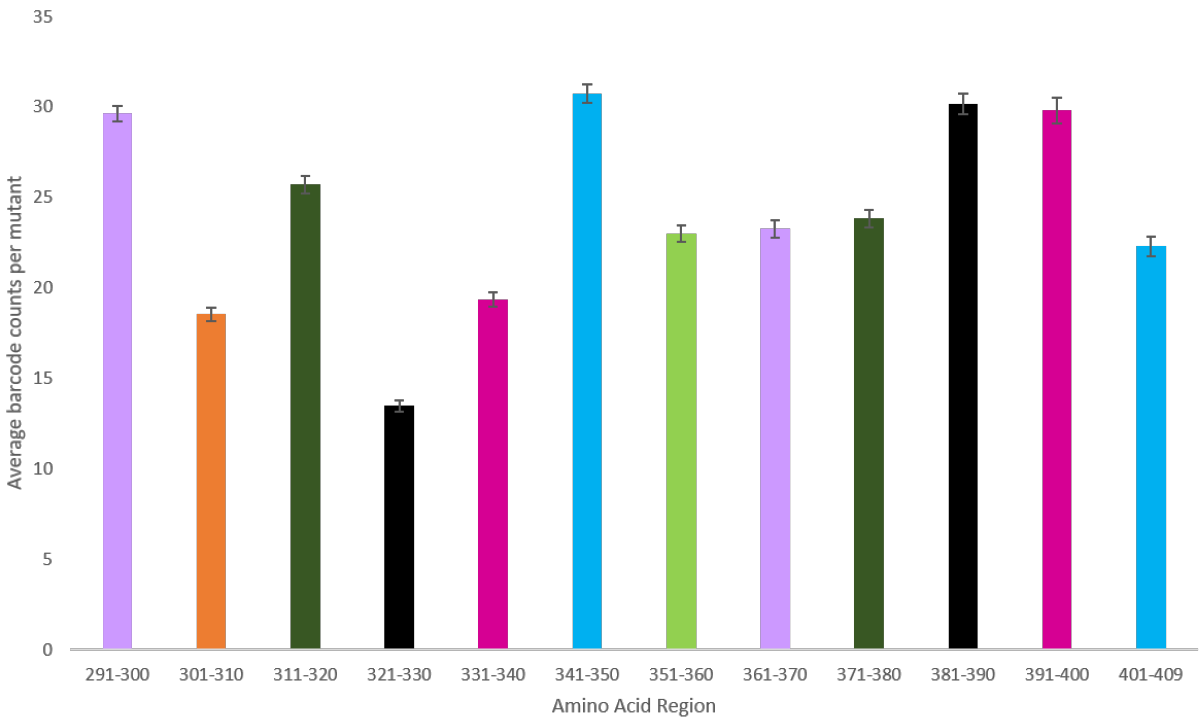


**Supplementary figure 13**. Average number of barcodes per mutant for each 9 amino acid window shows an average of ~20 barcodes associated with each variant. The x-axis corresponds to groups of 10 amino acid regions and the y-axis to the average number of barcode counts per mutant.

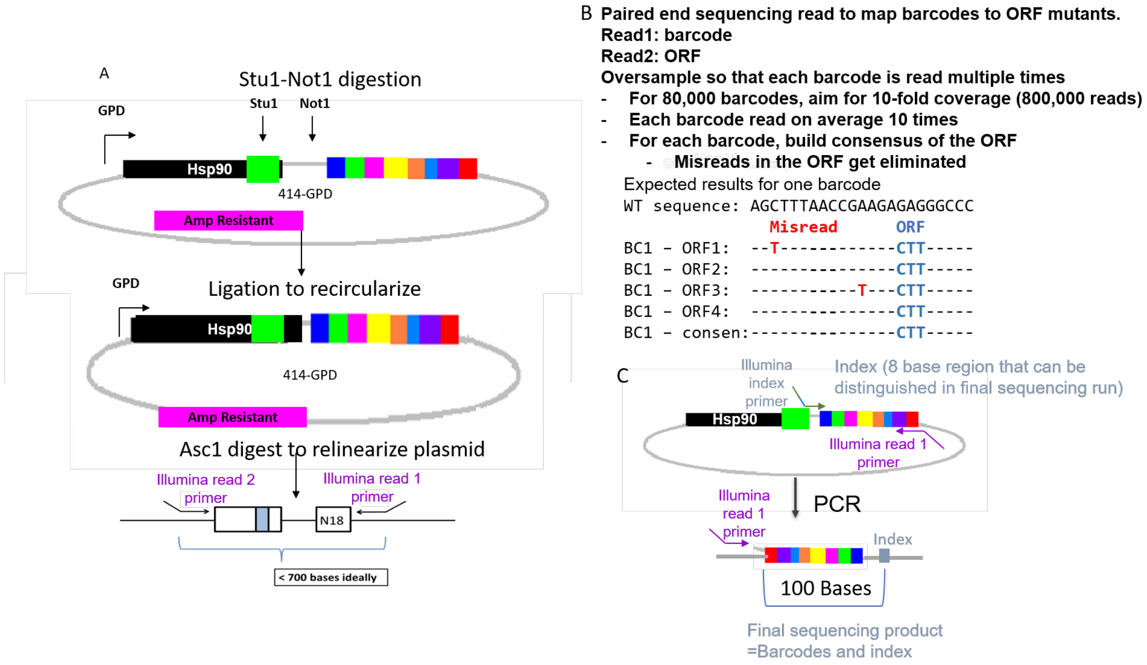


**Supplementary figure 14**. Strategy how to associate barcodes to mutants. A) Schematic of how to prepare barcoded mutant libraries for paired end deep sequencing. B) Schematic of how to map barcodes to open reading frame mutants. C) Schematic of how to include an index primer to distinguish each time point for final sequencing read following competition experiments.

### Supplementary Tables

#### Supplementary Table 1

**Number of beneficial, neutral and deleterious mutations across environments and replicates.** We observed a large proportion of wild-type like mutations in all environments.

| Environment | Replicate | % Beneficial | % Neutral | % Deleterious | Total Mutations |
|---|---|---|---|---|---|
| H202 | 1 | 5.03 (125) | 50.2 (1248) | 44.77 (1113) | 2486 |
| | 2 | 2.65 (65) | 54.5 (1337) | 42.85 (1051) | 2453 |
| Diamide | 1 | 12.34 (307) | 34.74 (864) | 52.92 (1316) | 2487 |
| | 2 | 10.73 (266) | 41.19 (1021) | 48.08 (1192) | 2479 |
| High Salinity | 1 | 2.57 (64) | 59.58 (1486) | 37.85 (944) | 2494 |
| | 2 | 4.85 (121) | 59.34 (1480) | 35.81 (893) | 2494 |
| Sorbitol | 1 | 7.09 (176) | 51.37 (1276) | 41.55 (1032) | 2484 |
| | 2 | 9.97 (245) | 60.09 (1477) | 29.94 (736) | 2458 |
| Standard | 1 | 9.03 (225) | 65.53 (1633) | 25.44 (634) | 2492 |
| | 2 | 6.14 (153) | 66.75 (1664) | 27.12 (676) | 2493 |
| | 3 | 5.19 (129) | 49.6 (1232) | 45.21 (1123) | 2484 |

#### Supplementary Table 2

**Pearson correlation of fitness effects across replicates and environments**. Blue indicates weaker correlations and red stronger correlations. Double lines indicate correlations between replicates of the same environment. In general the lowest correlations are seen for diamide and salt. Correlations between replicates are in general strong, except for standard replicate 3.

| | | Diamide | | H2O2 | | Sorbitol | | Salt | | Standard | | |
|---|---|---|---|---|---|---|---|---|---|---|---|---|
| | | Rep 1 | Rep 2 | Rep 1 | Rep 2 | Rep 1 | Rep 2 | Rep 1 | Rep 2 | Rep 1 | Rep 2 | Rep3 |
| Diamide | Rep 1 | | | | | | | | | | | |
| | Rep 2 | 0.96 | | | | | | | | | | |
| H2O2 | Rep 1 | 0.79 | 0.68 | | | | | | | | | |
| | Rep 2 | 0.78 | 0.70 | 0.96 | | | | | | | | |
| Sorbitol | Rep 1 | 0.76 | 0.66 | 0.95 | 0.94 | | | | | | | |
| | Rep 2 | 0.69 | 0.62 | 0.88 | 0.90 | 0.87 | | | | | | |
| Salt | Rep 1 | 0.64 | 0.48 | 0.86 | 0.78 | 0.84 | 0.73 | | | | | |
| | Rep 2 | 0.65 | 0.50 | 0.86 | 0.80 | 0.85 | 0.73 | 0.97 | | | | |
| Standard | Rep 1 | 0.63 | 0.47 | 0.86 | 0.77 | 0.82 | 0.71 | 0.96 | 0.95 | | | |
| | Rep 2 | 0.64 | 0.48 | 0.86 | 0.79 | 0.84 | 0.72 | 0.94 | 0.96 | 0.97 | | |
| | Rep3 | 0.81 | 0.71 | 0.99 | 0.97 | 0.95 | 0.90 | 0.83 | 0.83 | 0.82 | 0.83 | |

Correlation between replicates of the same environment

**Supplementary Table 3**

**Percentage (and number) of beneficial mutations and their cost across environments.** Beneficial mutations in a focal environment (rows) are deleterious in the environment indicated by the column. The diagonal of the matrix indicates the number of beneficial mutations found in each environment/replicate. Yellow shading indicates "spurious costs of adaptation" between pairs of replicates, which is a proxy for the expected classification error. In general, this error is low (between 0.38% and 6.61%), except for Replicate 3 in the standard environment and replicate 2 from Sorbitol, where many beneficial mutations in one replicate are classified as deleterious in the other.

| | | | Deleterious in | | | | | | | | | | |
|---|---|---|---|---|---|---|---|---|---|---|---|---|---|
| | | | Diamide | | Salt | | Standard | | | H2O2 | | Sorbitol | |
| | | | Rep1 | Rep2 | Rep1 | Rep2 | Rep1 | Rep2 | Rep3 | Rep1 | Rep2 | Rep1 | Rep2 |
| Beneficial in | Diamide | Rep1 | 307 | 0 (0) | 16.61 (51) | 18.24 (56) | 6.84 (21) | 8.14 (25) | 24.43 (75) | 26.38 (81) | 28.01 (86) | 22.8 (70) | 14.01 (43) |
| | | Rep2 | 0.38 (1) | 266 | 19.17 (51) | 18.05 (48) | 9.02 (24) | 9.02 (24) | 28.95 (77) | 30.83 (82) | 30.45 (81) | 25.19 (67) | 15.79 (42) |
| | Salt | Rep1 | 26.56 (17) | 25 (16) | 64 | 6.25 (4) | 0 (0) | 4.69 (3) | 1.56 (1) | 3.12 (2) | 9.38 (6) | 7.81 (5) | 1.56 (1) |
| | | Rep2 | 28.93 (35) | 23.14 (28) | 6.61 (8) | 121 | 4.13 (5) | 0.83 (1) | 9.09 (11) | 11.57 (14) | 11.57 (14) | 6.61 (8) | 0.83 (1) |
| | Standard | Rep1 | 28.89 (65) | 27.11 (61) | 7.56 (17) | 10.22 (23) | 225 | 2.22 (5) | 20.44 (46) | 20 (45) | 20.44 (46) | 19.11 (43) | 8.89 (20) |
| | | Rep2 | 33.99 (52) | 32.68 (50) | 9.15 (14) | 5.88 (9) | 3.92 (6) | 153 | 24.18 (37) | 24.18 (37) | 19.61 (30) | 12.42 (19) | 5.88 (9) |
| | | Rep3 | 22.48 (29) | 19.38 (25) | 4.65 (6) | 4.65 (6) | 2.33 (3) | 3.88 (5) | 129 | 0 (0) | 10.08 (13) | 2.33 (3) | 0 (0) |
| | H2O2 | Rep1 | 24 (30) | 17.6 (22) | 4.8 (6) | 3.2 (4) | 2.4 (3) | 4 (5) | 0.8 (1) | 125 | 4.8 (6) | 6.4 (8) | 1.6 (2) |
| | | Rep2 | 21.54 (14) | 16.92 (11) | 4.62 (3) | 1.54 (1) | 6.15 (4) | 1.54 (1) | 4.62 (3) | 0 (0) | 65 | 3.08 (2) | 1.54 (1) |
| | Sorbitol | Rep1 | 27.84 (49) | 23.3 (41) | 5.68 (10) | 2.27 (4) | 2.27 (4) | 2.84 (5) | 5.11 (9) | 7.39 (13) | 10.23 (18) | 176 | 3.98 (7) |
| | | Rep2 | 34.29 (84) | 29.8 (73) | 7.35 (18) | 3.67 (9) | 4.9 (12) | 4.08 (10) | 12.24 (30) | 16.73 (41) | 13.06 (32) | 15.1 (37) | 245 |

| | |
|---|---|
| (yellow) | Percentage of beneficials that have deleterious effect between replicates |
| (grey) | Number of beneficials in each environment |
